# Where did you come from, where did you go: Refining Metagenomic Analysis Tools for HGT characterisation

**DOI:** 10.1101/401349

**Authors:** Enrico Seiler, Kathrin Trappe, Bernhard Y. Renard

## Abstract

Horizontal gene transfer (HGT) has changed the way we regard evolution. Instead of waiting for the next generation to establish new traits, especially bacteria are able to take a shortcut via HGT that enables them to pass on genes from one individual to another, even across species boundaries. Existing HGT detection approaches usually first identify genes of foreign nature, e.g., using composition-based methods, and then exploit phylogenetic discrepancies of the corresponding gene tree compared to a species tree. These approaches depend on fully sequenced HGT organisms and computable phylogenetic species trees. The tool Daisy offers a different approach based on read mapping that provides complementary evidence compared to existing methods at the cost of relying on the acceptor and donor references of the HGT organism being known. Acceptor and donor identification is akin to species identification in metagenomic samples based on sequencing reads, a problem addressed by metagenomic profiling tools. However, acceptor and donor references have certain properties such that these methods can not be directly applied. We propose DaisyGPS, a mapping-based pipeline that is able to identify acceptor and donor candidates of an HGT organism based on sequencing reads. To do that, DaisyGPS leverages metagenomic profiling strategies and refines them for HGT candidate identification. These candidates can then be further evaluated by tools like Daisy to establish HGT regions. We successfully validated our approach on both simulated and real data, and show its benefits in an investigation of MRSA outbreak data. DaisyGPS is freely available from https://gitlab.com/rki_bioinformatics/.

## 1 Introduction

For a long time, evolution in terms of gene transfer was thought to happen only along the tree of life, i.e. from parent to offspring generation. The discovery of horizontal gene transfer (HGT) (Ochman et al., 2005, Boto, 2009, Wiedenbeck and Cohan, 2011, Daubin and Szöllősi, 2016) has revolutionised this dogma, and revealed the mechanism that enables bacteria to quickly adapt to environmental pressure (Hu et al., 2011, McElroy et al., 2014, Gyles and Boerlin, 2013). Via HGT, bacteria can directly transfer one or multiple genes from one individual to another across species boundaries. The known and prominent mechanisms of HGT are transformation (uptake of nascent DNA from the environment), conjugation (direct transfer from cell to cell), and transduction (transfer via bacteriophages) (Gyles and Boerlin, 2013). In all cases, a piece of DNA sequence is-directly or indirectly-transferred from the so called donor organism to the acceptor organism and integrated into the genome (see also Figure 1). Especially conjugation and transduction facilitate the transfer of pathogenicity islands and mobile genetic elements involving antimicrobial resistance (AMR) genes (Barlow, 2009, Warnes et al., 2012, Juhas, 2013). Today, we are facing the rise of so called ”superbugs” (Juhas, 2013, Perry et al., 2014) as a result of bacterial adaptation and gain of resistance to antibiotic treatment, showing the need for methods to identify, characterise and trace HGT events.

**Figure 1:**
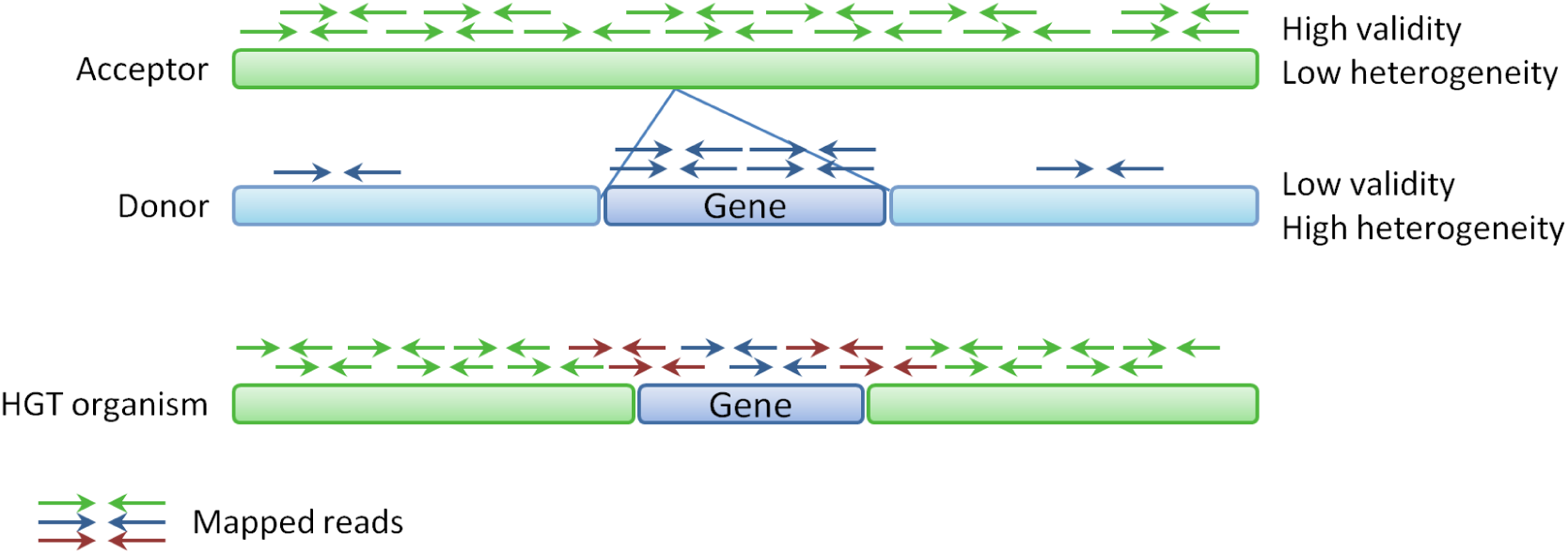
HGT overview and evidence. The sequence of an HGT organism consists mainly of the sequence of the acceptor genome (green), and only the transferred part (blue gene) is represented by the donor genome. Hence, reads from the HGT organism should mainly map homogeneously to the acceptor (green arrows), only few reads should map locally to the donor (blue arrows), and some read pairs (red arrows) will span the boundary between the green parts from the acceptor and the blue part from the donor. These mapping patterns can be represented by scores based on the mapping coverage profile. An acceptor with a homogeneous coverage has a high validity score and a low heterogeneity score, a donor has opposite score ranges (low validity and high heterogeneity). Based on these scores, the DaisyGPS *acceptor*-*score* is ∈ [0, 1] and *donor*-*score* is ∈ [-1, 0).

The discrepancy to phylogenetic evolution inspired existing genome-based HGT methods. For a fixed set of species and a potential horizontally transferred gene, these methods detect HGT events by looking at inconsistencies between the gene tree and a phylogenetic tree built for the set of species (Ravenhall et al. (2015)). As a prerequisite, a candidate gene for which to run the calculation and comparison has to be known. Sequence content based methods aim to identify genes of foreign origin in a given genome by exploiting sequence pattern such as *k-mer* frequencies or GC content which vary between different species (Jaron et al. (2013), Metzler and Kalinina (2014)). All methods are based on an assembled HGT organism, meaning they are also prone to the problems of misassemblies. Although AMRs are a prominent example for horizontally transferred genes, methods to directly identify antimicrobial resistance (AMR) genes do not necessarily connect the presence of an AMR gene to an HGT event (e.g., KmerResistance Clausen et al. (2016)).

In previous work, we developed an approach that aims to call HGT events directly from next-generation sequencing (NGS) data (Trappe et al., 2016) in a tool called Daisy. Instead of focusing on the sequence content of the HGT organism, Daisy examines the origin of the transfer, namely the prespecified acceptor and the donor organisms, and directly maps the NGS reads to these references. By facilitating structural variant detection methods, we can thereby identify the transferred region from the donor and the insertion site within the acceptor. A prerequisite for Daisy is therefore that both acceptor and donor references are known. This, however, is not always the case, and hence requires methods that are able to infer acceptor and donor candidates from the NGS reads of the HGT organism. Such methods are not yet available.

However, the problem of acceptor and donor identification directly from NGS data of the HGT organism is akin to the problem tackled by metagenomic profiling studies that aim to unravel metagenomic samples. Here, so called metagenomic classification approaches aim at identifying all organisms present in a sample by directly analysing sequencing data with a complex mixture of various organisms (Breitwieser et al., 2017). While in this classical scenario all reads of a single organism in the sample can theoretically be assigned to one reference organism during identification, this is not the case for an organism that carries foreign genes acquired via HGT. Most reads will be assigned to the acceptor genome but only a fraction can map to the donor genome (see mapped reads in Figure 1). Hence, we have to account for this two mapping properties of the reads during analysis. Another requirement is the resolution of classification on strain level, if possible, since two strains of the same species can already significantly differ in their sequence content.

Metagenomic classification approaches follow either a taxonomy dependent or taxonomy independent approach (Lindgreen et al. (2016), Sedlar et al. (2017)). The general procedure for both approaches is to assign sequencing reads stemming from the same organism in the sample into the same group, a process also referred to as binning. Taxonomic dependent binning approaches assign the reads to specific taxonomic groups, and hereby infer the presence of these taxa in the sample. These methods either also make use of sequence composition patterns, e.g., Kraken (Wood and Salzberg, 2014), or they determine mapping-based sequence similarities for the read assignment, e.g., MEGAN (Huson et al., 2007), Clinical PathoScope (Byrd et al., 2014) or DUDes (Piro et al., 2016). Both approaches will most likely identify the acceptor reference of an HGT organism due to the homogeneous coverage and comparatively high number of reads. The drawback of all read assignment approaches is the limitation in the presence of mobile genetic elements, e.g., integrated via HGT or of hitherto unknown-or un-sequenced-organisms in the sample. Reads belonging to these genes or unknown organisms are either assigned to a similar but incorrect taxa or not assigned at all, leading to wrong identifications and biases in abundance estimation. To ensure robustness, many approaches deliberately discard taxonomic candidates with only low and local coverage. Hence these approaches will likely discard any donor candidate references. Composition-based methods such as Kraken would also perform poorly pin-pointing the correct donor based on evidence of only few reads given the fairly large number of usually detected species.

In our group, we developed MicrobeGPS (Lindner and Renard, 2015), a metagenomics approach that accounts for sequences not yet present in the database. Instead of reporting fixed taxa with assigned reads, MicrobeGPS in turn uses the candidate taxa to describe the organisms in the sample in terms of a genomic distance measure. That is, it uses available references to model the composition of the organisms present in the sample in terms of coverage profiles and continuity, instead of directly assigning reference organisms to characterize the sample. If the organism in the sample is present in the database and covered homogeneously then the distance approximates to zero. If not, MicrobeGPS identifies the closest relatives by positioning the organism among references with the lowest genomic distance. Hence, the tool considers scores and metrics that reflect a donor-like, in-homogeneous coverage but filters out false positive candidates with inhomogeneous coverage for the purpose of species assignment. From the perspective of HGT detection, these may be highly relevant and should not be excluded.

Here we present DaisyGPS, a pipeline building on concepts of MicrobeGPS and tailored to the identification of acceptor and donor candidates from sequencing reads of an HGT organism. DaisyGPS uses genome distance metrics to define a score that allows the classification into acceptor and donor among the reported organisms. Owing to the properties of these scores, we still find the closest relatives of acceptor and donor in case these references are not present in the database. DaisyGPS further offers optional blacklists and a species filter to refine the search space for acceptor and donor candidates. DaisyGPS and Daisy are integrated into one pipeline called DaisySuite to offer a comprehensive HGT detection, and publically available at https://gitlab.com/rki_bioinformatics/DaisySuite. We validate Daisy-Suite on a large scale simulation where we show sensitivity and specificity of our approach and the robustness when applied to non-HGT samples. On a real data set from an MRSA outbreak, we demonstrate the ability of the DaisySuite to distinguish between the outbreak associated and unassociated samples in terms of sequenced content potentially acquired through HGT events.

## 2 Methods

The problem of mapping-based HGT detection from NGS data is twofold: First, the acceptor (organism that receives genetic information) and donor (organism that the information is transferred from) references have to be identified. Based on that, the precise HGT region and its insertion site within the acceptor can be characterised. We presented a method to solve the second task in Trappe et al. (2016). Here, we propose the tool DaisyGPS (see also Figure 2) with the objective to identify possible acceptor and donor candidates given reads of a potential HGT organism. We provide Daisy and DaisyGPS in an integrated pipeline that we call Daisy-Suite.

**Figure 2:**
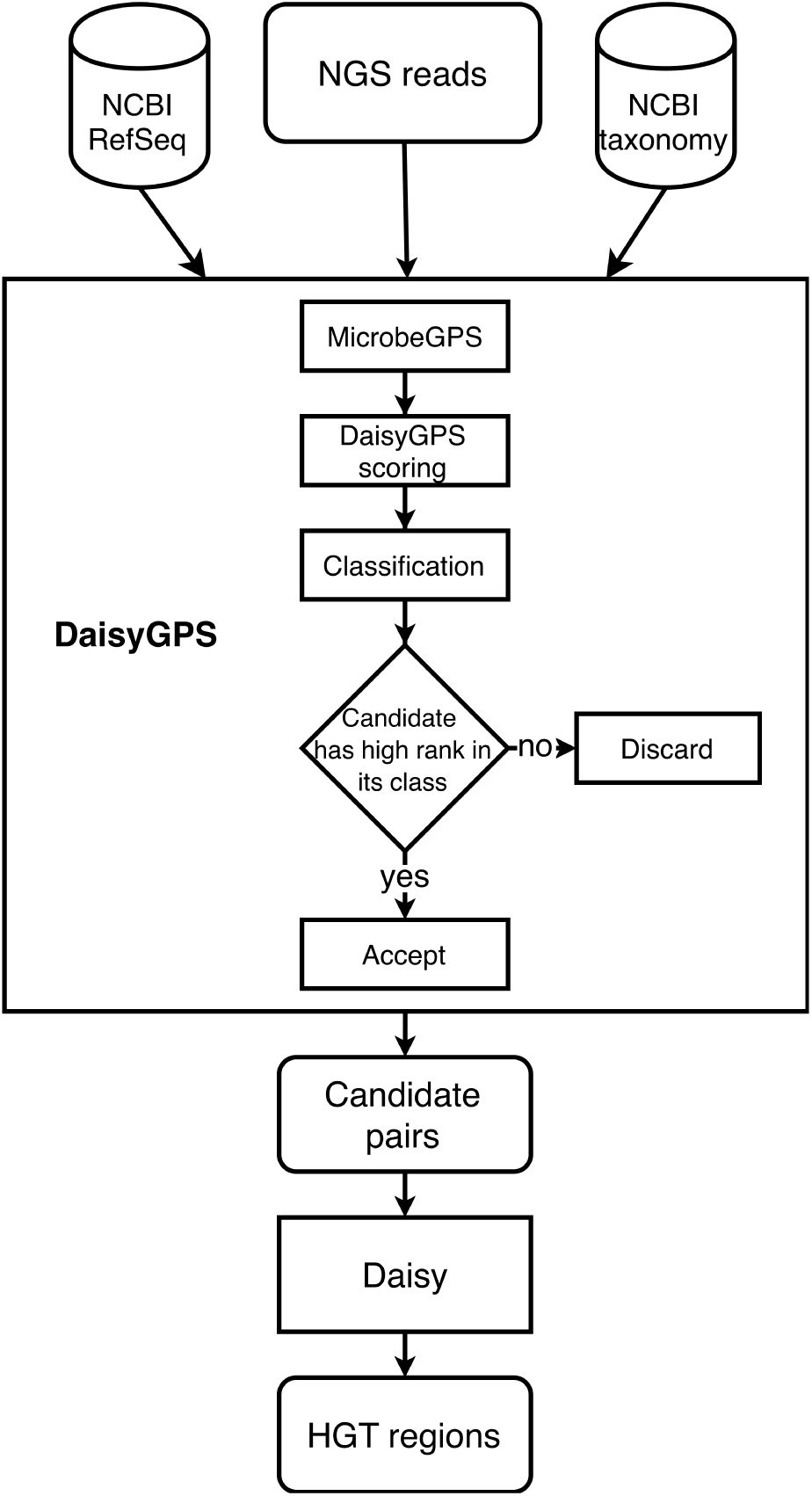
Workflow of DaisySuite. The input NGS reads are first processed by DaisyGPS. The reads are mapped to the NCBI RefSeq and then analysed by MicrobeGPS which also incorporates taxonomic information acquired through the NCBI taxonomy database. Based on that, DaisyGPS calculates two scores for acceptor and donor classification (see methods part). Depending on these scores, the highest-ranked candidates are selected as suitable acceptor and donor candidates. Daisy then uses these candidates to identify HGT region candidates.

**Figure 3:**
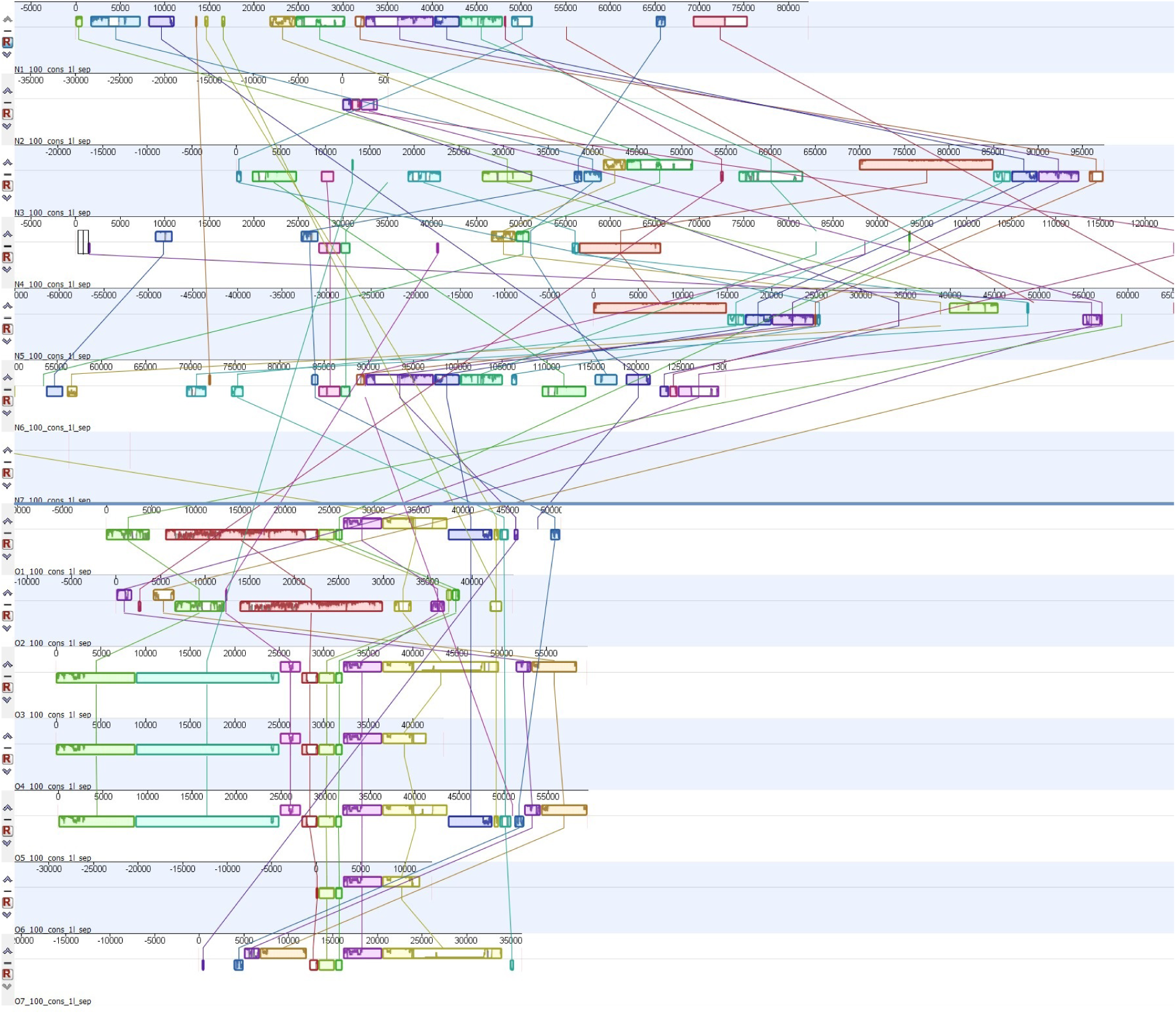
Mauve alignment of concatenated HGT regions. The HGT regions of all samples are aligned with Mauve to establish shared regions between them. The outbreak associated samples (O1-O7) in the lower part share most of their regions whereas the unassociated samples (N1-N7) in the upper part do not.

The genome of the HGT organism consists mainly of the acceptor genome (see Figure 1). When the reads of the HGT organism are mapped against the acceptor reference, most reads should map properly. Therefore a high and continuous mapping coverage pattern of the acceptor genome can be expected. In contrast to that, only a small part of the donor genome is present within the genome of the HGT organism, hence only a small fraction of the reads should map against the donor reference and then only within a zoned part (i.e. the part that has been transferred). This results in a discontinuous mapping coverage pattern where only a small part of the reference shows a high mapping coverage (see Figure 1).

In a first step, we need to define metrics that represent the expectations we have, i.e. how much of the genome is covered by reads (mapping coverage) and how uniformly these reads are distributed across the genome (discontinuous vs. continuous patterns). Given only the reads of the HGT organism, the acceptor and donor candidate identification problem is similar to aspects of metagenomic profiling. A standard problem in metagenomics is the identification of organisms in a sample using a read dataset of this sample. At first glance, it may appear that the methods designed to solve this problem can also be applied to our identification objective, i.e. we have the read dataset of the HGT organism and we are looking for two organisms (acceptor and donor) that are in the sample. However, because the HGT organism consists mainly of the acceptor genome, such an approach works only well for the identification of the acceptor. For the donor, additional information is needed to guarantee a reliable identification because references with only local or discontinuous coverage are usually dismissed by the profiler. We use the metagenomic profiling tool MicrobeGPS to obtain a coverage profile of our given HGT organism from mapping coverage metrics. MicrobeGPS fits our requirements as it can be configured to not filter any organisms and reports additional metrics that we use to represent acceptor and donor attributes. Next, we evaluate the gathered metrics and establish a score that reflects our defined acceptor or donor coverage properties. Then, the candidates are ranked by this score and a list of acceptor and donor candidates is generated. These acceptor and donor candidates can then be further analysed with tools such as Daisy.

### DaisyGPS scores

For the purpose of HGT detection, we aim to define a scoring that reflects the mapping coverage properties of the acceptor and donor references: The acceptor has a continuous, homogeneous coverage over the complete length of the genome. The donor has a local, but still homogeneous coverage in the area where the transferred genes are originated but should have nearly no coverage at all otherwise. The score should further allow a clear distinction between acceptor and donor candidates and provide a meaningful ranking according to the likelihood of being the most suitable candidate.

As a basis for our scoring, we use the *Genome Dataset Validity* defined in Lindner et al. (2013) and *homogeneity* metric defined in Lindner and Renard (2015). The Genome Dataset Validity, or short validity, describes ∈the fraction of the reference genome for which there is read evidence. In contrast, the homogeneity reflects how evenly the reads are distributed. Both have a range ∈ [0, 1]. The validity is defined such that a genome that is covered-either low or high-over the full length has a high validity (≈1). We define a *heterogeneity* metric based on the Kolmogorov-Smirnov test statistic defined in Lindner and Renard (2015) such that an evenly covered genome has a low heterogeneity (≈ 0) and a genome with local, high coverage a high heterogeneity (≈ 1).

An acceptor is a genome with a continuous, high coverage that then has a high validity (≈1) and a low heterogeneity (≈0) score whereas a distantly related donor genome with only local, discontinuous coverage has a low validity (≈ 0) and a high heterogeneity (≈ 1) score.

As can be seen above, both validity and heterogeneity are complementary for acceptors and donors, and hence the relation of both metrics infers the property of a candidate between being an acceptor or a donor candidate. We define: *score = validity* – *heterogeneity* with *score* ∈ [-1, 1]

Therefore, the value for a completely covered acceptor with uniform read distribution would approach +1. Likewise, the value for a donor that is only covered in a small region would approach-1. In addition to the coverage profile, there is a high evidence by sheer read numbers for acceptors:

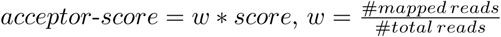

where *w* is the fraction of all mapped reads that mapped to the specific acceptor candidate. For the donor, however, the size of the transferred region is not known in advance. Hence, we do not expect a specific read number evidence and therefore omit the weighting and define *donor*-*score* = *score*

Both *acceptor*-*score* and *donor*-*score* are determined for every candidate and they have a codomain of [-1, 1]. Acceptor candidates have a homogeneous coverage and hence high validity and low heterogeneity, i.e. *validity* > *heterogeneity*. Hence, we classify the candidates with *acceptor*-*score* ≥0 as acceptor and rank them from highest to lowest score. Donor candidates have a high heterogeneity and low validity, i.e. *validity < heterogeneity*. Therefore, we classify candidates with *donor*-*score <* 0 as donor candidates and rank them from lowest to highest score.

There is a special case if acceptor and donor are very similar. Here, the donor might not express the attributes we are looking for. In particular, the donor might have a significant read number evidence arising from acceptor reads also mapping to the donor. These shared reads lead to more regions of the donor genome being covered (higher validity) and to a less local, more homogeneous coverage pattern across the donor genome (lower heterogeneity), hence *validity* ≈ *heterogeneity* and *donor*-*score* ≈ 0. We classify candidates with a *donor*-*score >* 0 as acceptor-like donors and rank them from lowest to highest.

### Candidate selection with blacklist filter (optional)

There are scenarios where it is necessary to exclude certain results from being reported. For example, in a reanalysis case, the assembled sequence from the sample reads might already been added to the reference set of your choice. For HGT detection from such reads, however, there is no information gain if DaisyGPS reports this entry as a suitable acceptor. Other examples include cases, where one can exclude certain species or taxa due to preanalysis information that nevertheless could be reported by DaisyGPS due to their high sequence similarity to the sampled organism or the presumed acceptor or donor candidates. To make the search for acceptor and donor candidates adaptable for such cases, DaisyGPS features the blacklisting of certain taxa. It is possible to exclude single taxa, a complete species taxon or a complete subtree below a specified taxon. For a default run, the filter is turned off.

### Candidate selection with species filter (optional)

DaisyGPS generally considers candidates on different taxonomic levels, e.g. species and strain level, and reports the candidate level with the best scores. Often the strain references contain additional sequences compared to the species level reference representative, and hence, the species reference will mostly have a homogeneous coverage that will then lead to a high acceptor score. Usually identification on species level is sufficient. There are however species such as, e.g., *E.coli*, where a high number of strains have been sequenced already and differ in their properties such as pathogenicity among the strains (e.g. *E.coli* K12 versus EHEC strain O157:H7). In these cases, a mere detection of the acceptor or donor on a species level might not be precise enough. For these situations, we implemented a species filter. If this filter is activated, only candidates below species level are reported. In case no candidate would be reported with an active species filter, the filter is disabled and the user informed that for further analysis also candidates on species level are used. For a default run, this filter is also turned off.

### Daisy inference and integration with Snakemake

Snakemake is a common workflow management system (Köster and Rahmann, 2012) which we used to implement the different steps of DaisyGPS. We generated the alignment file required for MicrobeGPS by mapping the reads of the HGT organism against the NCBI RefSeq (complete RefSeq, no plasmids, downloaded March 15th 2017) (O’Leary et al., 2016) using Yara (Siragusa et al., 2013, Dadi et al., 2018). To ensure compatibility, we reimplemented the Daisy workflow in Snakemake as well, and integrated both into a combined suite (called DaisySuite, see also Figure 2). DaisyGPS yields a configurable number of acceptors, donors and acceptor-like donors (default: 2, 3, 2). For each possible pair of acceptor and donor, a Daisy call is inferred. Both pipelines can still be run independently. To unburden installation, we provide a setup script and provide DaisySuite components as Conda (Con) packages. The simulations are also integrated into the DaisySuite pipeline (see DaisySuite documentation for details).

## Experimental setup

### Data sets

We tested the complete DaisySuite on three types of data sets to validate both DaisyGPS and the integration with Daisy. The first type comprises the *H.pylori* data set, the KO11FL data set and the EHEC data set. All three were used in the Daisy publication (see Trappe et al. (2016) for detailed data set description) and are chosen as suitable ground truth and for the purpose of showing reproducibility. The second type comprises a large-scale simulation analogous to the *H.pylori* simulation. Both positive (simulated HGT) and negative (no HGT) simulations are used to estimate sensitivity and specificity of the DaisySuite. In a third part, we use real data from an outbreak data set with 14 MRSA samples to elucidate further applicability of both DaisySuite. The details of the data sets and *in silico* experiments are explained below.

#### H. pylori

The data set *Helicobacter pylori* presents a simulated data set for a proof of principle already used for validation in the Daisy paper (see Trappe et al. (2016) for details of genomic simulation). The acceptor is *Escherichia coli* K12 substr. DH10B (NC 010473.1), the donor is *H. pylori* strain M1 (NZ AP014710.1). The *in silico* transferred phage region of the *H. pylori* comprises genomic positions 1 322 000-1 350 000.

#### EHEC

The HGT organism in the EHEC data set is *E.coli* O157:H7 Sakai (Zhang et al., 2007) that derived from *E.coli* O55:H7 and is assumed to have acquired the Shiga-Toxins (Stx) via transduction from *Shigella dysenteriae*. According to literature, the bacteriophage carrying Stx is supposedly positioned at 2 643 556-2 694 691 in *E.coli* O55:H7. In Trappe et al. (2016) we proposed an alternative phage insertion site at 1 741 535-1 744 926.

#### KO11FL

The KO11FL data set comprises the transgenic *E.coli* KO11FL (Turner et al., 2012). The acceptor is *E.coli* W, and the two donors are *Zymomonas mobilis* and the cloning vector pBEN77.

#### Large-scale simulation

We designed a large-scale simulation analogous to the *H.pylori* data set with positive and negative simulations. For each positive simulation, first an acceptor and a donor organism are randomly chosen among the available RefSeq sequences (date of retrieval: March 21, 2017, plasmids are ignored for sake of size consistency). A random 28 Kbp region is selected from the donor and inserted at a random position in the acceptor. SNPs and indels are introduced into acceptor and donor region (SNP rate: 0.01, indel rate: 0.001). For each negative simulation, only an acceptor is randomly chosen, and SNPs and indels are introduced with the same rates as above. 150 bp reads are simulated from 500 bp fragments with 50 bp standard deviation with the Mason simulator (Holtgrewe, 2014). The positive and negative simulations are repeated automatically 100 times.

#### MRSA outbreak

The MRSA data set consists of 14 samples of methicillin resistant *Staphylococcus aureus* strains obtained during a MRSA outbreak at a neonatal intensive care unit (ENA accession number ERP001256, Köser et al. (2012)). Seven samples are associated with the outbreak, labeled O1-O7 in this manuscript, the other seven samples N1-N7 are not associated with the outbreak. Sample description and run accession numbers are stated in Table 4. Phylogenetic analysis by Köser et al. (2012) separated the 14 samples into distinct groups according to their outbreak association. The reference isolate used in that study is the EMRSA-15 representative HO 5096 0412, and we use this as ground truth for acceptor candidates reported by DaisyGPS. The seven outbreak related MRSA samples have a distinct antimicrobial resistance pattern, and it is believed that the related resistance genes have been introduced via HGT. With DaisySuite we want to investigate if the outbreak strains share the same HGT regions and if they can be distinguished from the non-outbreak strains.

### Structure of validation

The setup of the validation is according to the types of data sets. In a first phase, we want to show a proof of concept given data with sufficient ground truth. The aim is to predict the correct acceptor and donor candidates with DaisyGPS and at the same time to reproduce the results obtained from Daisy. We therefore use the data sets already shown in the Daisy paper for sake of consistency. We set DaisyGPS to report a total of two acceptor candidates, four donor candidates, and two acceptor-like donor candidates for every data set and we evaluate if the correct acceptor and donor candidates are among them. For incorrect candidates of acceptor and donor, Daisy should not report HGT candidates unless the transferred region is present in multiple strains or there are multiple possible acceptors present with high sequence similarities as, e.g., among *E.coli* strains. For the EHEC data set, we activate the species filter since we are interested in strain candidates, and further blacklist taxa from the HGT organism to be analysed (*E.coli* O157:H7, taxon 83334) and the complete O157 lineage (parent taxon 1045010). For the KOFL11 data set, the HGT organism is blacklisted as well (*E.coli* KOFL11, taxon 595495). In a second part, we want to estimate the rate of sensitivity and specificity of the DaisySuite. We designed a large-scale simulation analogous to the *H.pylori* data set with positive and negative simulations (100 simulations each). From the positive simulations, we calculate the sensitivity for both DaisyGPS and Daisy (see below for definitions on metrics). DaisyGPS is designed with high sensitivity in mind and always reports the closest fitting candidates given sequencing data, even for non-HGT organisms. Hence, also for the negative simulations, DaisyGPS will report candidates and we expect a low specificity here. Daisy, however, should then report only few if any-HGT candidates from the acceptor-donor pairs. In the last evaluation part, we test the DaisySuite on real data with unknown or uncertain ground truth. The MRSA outbreak data set consists of 14 samples, seven outbreak related and seven unrelated. Here we want to test if DaisySuite is able to distinguish between the outbreak and non-outbreak samples according to their reported acceptor, donor and HGT region candidates.

### Definition of evaluation metrics

The interpretation of various statistics depends on the hypothesis to be tested. In our analysis in the large-scale simulations, we differentiate between two scenarios: in the first one we expect to detect an HGT event (positive test), while in the other one we assume the absence of an HGT event (negative test). For each simulation or run, a DaisyGPS call will lead to multiple pairs to be evaluated by Daisy. We therefore distinguish between statistics on runs and statistics on pairs that we will explain in the following.

For DaisyGPS, we consider during a positive test a single run as a true positive (TP) if the correct acceptor/donor pair is reported. Accordingly, a false negative (FN) occurs when the correct pair is not reported. Since the number of reported pairs is set by our settings, we will almost always have a fixed number of downstream verifications (except if there are not enough candidates to report) and thus we report the number of runs instead of pairs. Consequently, we can define the sensitivity as TP / #Runs. In a negative test setting, we deem those runs as true negatives (TNs) where either no pairs are reported or acceptor and donor of the pair are the very same organism. All other pairs are regarded as FP that will each trigger an unnecessary verification in the down-stream tools. Since we are interested in how many runs did not cause verifications, we can characterize the specificity by TN / #Runs. While it is obvious in both settings to rely on an exact match of the reported results and the ground truth, a reported organism still may be very close to the ground truth organism in terms of sequence similarity (negative and positive settings) and even include the very regions involved in the HGT event (positive setting). To account for this, we also use BLAST in the case that no TP was reported and compare the FP to the ground truth. If the Blast identity of the FP to the ground truth is above 80% we change the classification from FP to BLAST-supported TP (Blast TP) since Daisy might still be able to infer the correct HGT region from these Blast TPs given the sufficient sequence similarity.

In Daisy, we evaluate acceptor/donor pairs and therefore the statics are defined based on the condition of a pair reported by DaisyGPS. In a positive simulation, Daisy TP pairs are those that represent the correct pair and are detected by Daisy. It directly follows that each correct pair that is not supported by Daisy can be seen as a false negative (FN). Given that the pair is incorrect, i.e. a FP from DaisyGPS where the acceptor or donor is wrong, we count a rightly not supported pair as true negative (TN) and an erroneously detected pair as FP. To measure how many pairs are correctly identified, we define the sensitivity as (TP + TN) / #Pairs. Considering a negative test setting, we are mainly interested in the pairs that are wrongly reported as being involved in an HGT event. We declare those pairs as FP and describe the specificity as (#Pairs-FP) / #Pairs. It also follows that all the pairs that are not detected are TN.

Lastly, in the context of the complete DaisySuite pipeline, we evaluate the combined results of DaisyGPS and Daisy. Each pair reported by DaisyGPS for a single simulation induces an evaluation by Daisy. Since the overall result of the pipeline should indicate whether a simulation contains an HGT event or not, the classification of a DaisySuite run depends exclusively on the consolidated results of each Daisy evaluation for a single simulation. In a positive test setting, we want to find exactly the one pair that represents the HGT event. From that follows that a complete DaisySuite run can be classified as TP if Daisy supports solely the correct pair, i.e. Daisy reports the TP and no FP. This also implies that DaisyGPS needs to detect the TP. Similarly, in a negative test setting, a TN occurs if Daisy reports no HGT candidates at all.

### Settings and pre-/post-processing

DaisySuite is run with default parameters as of version 0.0.1 unless stated otherwise. The parameter to combine potentially overlapping HGT candidates within Daisy is set to 20 bp, hence, overlapping regions with start and end positions differing by more than 20 bp are reported as separate candidates. For the comparison of the number and content of HGT sequences, we clustered overlapping HGT candidates with the tool usearch9 (v9.1.13 i86linux32) with identity 1.0 (Edgar, 2010).

For validation, we determine the true presence of a HGT region in the samples by mapping the sample reads to all suggested, clustered regions with Bowtie2 (version 2.2.4). For comparison, we take the mean coverage of every region and apply a sigmoidal function to map all mean coverages to the [0.5,1] space for displaying a meaningful heatmap. The application of a sigmoidal function and the heatmap is computed in R (Rscript version 3.3.3). The heatmap function in R uses a hierarchical clustering with complete linkage as default, and we turned of the dendrogram for the columns. In addition, we perform a whole-genome alignment using the Mauve plugin (version 2.3.1) as part of the Geneious software (version 10.0.5) to to establish shared HGT regions among the samples. To do this, we concatenate all HGT regions of a sample and separate the regions with segments of 1000*N to avoid fragmented regions or overlapping LCBs.

## 3 Results

### Acceptor and donor identification with DaisyGPS

In the first part of the validation, we test DaisyGPS on three data sets from simulated and real data with sufficient ground truth and already previously evaluated with Daisy. Since DaisySuite combines both tools, DaisyGPS and Daisy, the aim is to support our previous results even when now the donor and acceptor are not prespecified.

The *H.pylori* data set was simulated from *E.coli* K12 substr. DH10B as acceptor and *H. pylori* strain M1 as donor. DaisyGPS successfully reports both as such (see Supplement Tables S3 and S4), and the subsequent Daisy run also reports the true HGT site. In addition to the only true HGT candidate previously already reported in the Daisy paper, DaisySuite reports another, FP HGT site for a region from *Haemophilus ducreyi*. The HGT region reported for *H. ducreyi* strain GHA9 has no continuous similarity with the HGT region from *H.pylori* (no blast hits longer than 15 bp, data not shown). However, the region on *H. ducreyi* shares the first 1200 bp and the last 1300 bp with the acceptor *E.coli* K12 substr. DH10B on multiple sites, and since beginning and end of the region are covered, almost six times as many split-reads are found as for the true acceptor site. The total coverage of the region is relatively low with 30x compared to 95x of the *H.pylori* but obviously high enough to pass the coverage filter.

The EHEC *E.coli* O157:H7 Sakai is supposedly derived by an HGT event where a defective prophage has been transferred from *Shigella dysenteriae* to *E.coli* O55:H7. Both are reported by DaisyGPS as candidates (see Supplement Table S5). In line with its strong sequence similarity to the *E.coli* species, *S.dysenteriae* is labeled as an acceptor-like donor candidate. The proposed alternative HGT insertion site from our previous Daisy paper is still reported (see Supplement Table S6).

The KO11FL data set comprises a transgenic *E.coli* W variant with transferred genes from *Zymomonas mobilis* and a plasmid that was not analysed here. DaisyGPS successfully reports *E.coli* W and *Zymomonas mobilis* as acceptor and donor candidates (see Supplement Table S7). Daisy does not report any FP HGT candidates.

### Estimating sensitivity, specificity and robustness of DaisySuite through large-scale simulations

After validating DaisyGPS on data previously evaluated with Daisy as a proof of principle, we analyse DaisySuite in terms of robustness and sensitivity by performing a large-scale simulation. We perform the simulation for the *H.pylori* data set in a randomised and automated fashion generating 100 simulations with a transferred HGT region. To evaluate robustness, we also perform 100 negative simulations where an acceptor genome is simulated but no HGT region is inserted. With the positive simulations, we can estimate the sensitivity of the complete DaisySuite. For DaisyGPS, we evaluate how many from the 100 simulations have the correct acceptor and donor genome identified. Since DaisyGPS reports more than one potential acceptor-donor pair, we count a TP hit if the true pair is among them, and only count a FN if the true pair was not reported at all. In addition, we consider pairs with Blast sequence identity *>* 80% also as a potential HGT candidate pair, and also count them as a TP. To evaluate Daisy, we consider all pairs proposed by DaisyGPS.

For a true pair reported by DaisyGPS, Daisy can either report a TP HGT region or a FN if the region could not be identified. For an acceptor-donor pair wrongly proposed by DaisyGPS, Daisy can either report no HGT candidate region (TN) or a FP hit. When we summarise the DaisySuite results over all pairs of one simulation, we only count a TP for that simulation if Daisy did not report any FPs (despite any TPs or TNs).

Table 1 states the resulting counts for DaisyGPS and for the complete DaisySuite summarised over the 100 simulations. DaisyGPS yields a sensitivity of 79%. From the 79 TPs, 22 are based on either a wrong acceptor, or donor, or both but have still sufficient Blast similarity to the original acceptor or donor to be counted as TP according to our scoring. 69% of the TPs and FPs resulted in a TP or TN call from Daisy. It is noticeable that all DaisySuite FPs are Blast FPs.

**Table 1:**
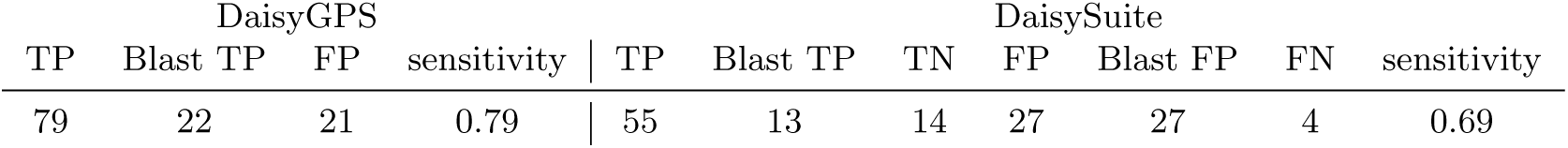
Positive HGT simulation. DaisyGPS calls correct acceptor and donor candidates with a sensitivity of 79%. The total sensitivity for DaisySuite from 100 HGT simulations regarding correct acceptor and donor candidates with a follow up correct HGT site call is 69%.

Table 2 states the number of reported pairs proposed by DaisyGPS and a detailed count based on each pair for Daisy. From the resulting 818 pairs, Daisy then reports the correct HGT region, or correctly no HGT region from a DaisyGPS FPs, with a sensitivity of 89%.

**Table 2:**
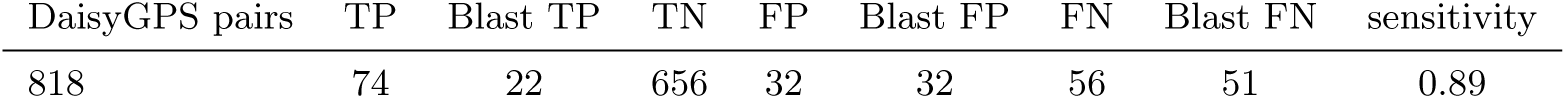
Positive HGT simulation. Daisy evaluates 818 pairs reported by DaisyGPS and calls the correct HGT region or correctly no HGT region with a sensitivity of 89%.

In addition to the positive simulations, we performed another 100 negative simulations where we randomly selected and variated an acceptor genome but did not insert any foreign region from a donor. DaisyGPS can now either produce a TN hit, i.e. report no candidates at all, or FP candidates. Since DaisyGPS is very sensitive by design, we expect it to report candidates most of the time and, hence, we want to estimate if these negative HGTs trigger reports by a Daisy follow-up call. As expected, the specificity for DaisyGPS is very low with 6% (see Table 3). However, Daisy reports only six FPs on all pairs in total, i.e. three simulations produced a FP HGT report.

**Table 3:** Negative HGT simulation. For the 100 negative simulations, DaisyGPS correctly reports no acceptor and donor candidates for six simulations. From the 94 simulations causing a downstream evaluation with Daisy, only three lead to a FP call considering all outcomes from DaisySuite (summarised over the 100 simulations). Daisy evaluates 743 pairs and only has six FP HGT region calls in total over all those pairs.

| DaisyGPS |  | DaisySuite |  | DaisyGPS pairs |  | Daisy |  |
| --- | --- | --- | --- | --- | --- | --- | --- |
| TN | specificity | FP | specificity |  |  | FP | specificity |
| 6 | 0.06 | 3 | 0.97 | 743 |  | 6 | 0.99 |

From these results we can infer that DaisySuite is able to distinguish HGT from non-HGT organisms and is very robust if no HGT is present.

### Exploration of HGT detection with DaisySuite from MRSA outbreak data

MRSA strains are generally assumed to undergo HGT events frequently (Lindsay, 2010, 2014). The MRSA data set considered here consists of 14 samples with seven of them related to an MRSA outbreak (O1-O7) and seven MRSA samples not associated with the outbreak (N1-N7) but that occurred in the same time frame (Köser et al., 2012). Köser et al. (2012) analysed all 14 samples and compared them to the EMRSA-15 representative HO 5096 0412 as the supposedly closest relative of the outbreak strains. We first evaluate acceptor and donor candidates reported by DaisyGPS in relation to the proposed HO 5096 0412 reference and then investigate HGT region candidates reported by Daisy regarding a possible distinction of outbreak vs. non-outbreak samples. We activate the species filter as we are again interested in strain level candidates.

For all outbreak samples O1-O7, *S.aureus* HO 5096 0412 was reported as acceptor candidate by DaisyGPS (see Table 4 and supplementary tables S8-S35). The same acceptor was also reported for non-outbreak samples N2, N6 and N7. Acceptor candidates for sample N1 are *S.aureus* ECT-R-2 and N315, for N3 and N4 *S.aureus* MSSA476 and MW2, and for N5 *S.aureus* MRSA252. Although not associated with the outbreak, samples N3 and N4 are from patients that shared the same room in the hospital where the outbreak occurred and hence are possibly related (Köser et al., 2012).

**Table 4:** Acceptor and number of HGT region candidates. For 10 of the 14 samples, EMRSA-15 (HO 5096 0412) was reported as acceptor candidate. This includes all outbreak samples. Column *HGT regions* states the number of reported HGT regions, and column *EMRSA-15 as acceptor for HGT regions* the respective number that were reported with HO 5096 0412 as acceptor.

| Label | Isolate | Accession | EMRSA-15 as acceptor | HGT regions | EMRSA-15 as acceptor for HGT regions |
| --- | --- | --- | --- | --- | --- |
| O1 | 1B | ERR103401 | x | 4 | 4 |
| O2 | 6C | ERR103403 | x | 4 | 3 |
| O3 | 7C | ERR103404 | x | 5 | 3 |
| O4 | 8C | ERR103405 | x | 3 | 3 |
| O5 | 10C | ERR101899 | x | 4 | 4 |
| O6 | 11C | ERR101900 | x | 1 | 1 |
| O7 | 12C | ERR103394 | x | 5 | 3 |
| N1 | 14C | ERR103395 | - | 5 | - |
| N2 | 15C | ERR103396 | x | 2 | 2 |
| N3 | 16B | ERR103397 | - | 4 | - |
| N4 | 17B | ERR103398 | - | 4 | - |
| N5 | 18B | ERR159680 | - | 5 | - |
| N6 | 19B | ERR103400 | x | 7 | 5 |
| N7 | 20B | ERR103402 | x | 2 | 2 |

The reported donors are largely the same for both outbreak and non-outbreak samples (see Table 5). No donor was reported exclusively for the outbreak samples but three donors only for non-outbreak strains N1, N4 and N6. These are *S.epidermidis* strains ATCC 12228 and PM221 as well as *Enterococcus faecium* Aus0004. Although *S.aureus* HO 5096 0412 was reported for all outbreak samples, there is no clear distinction in acceptor and donor candidates reported by DaisyGPS apart from the non-outbreak only donors.

**Table 5:** Reported donors summarised for all samples. Both outbreak associated and unassociated samples mostly report the same donor candidates with only few variations (see supplementary tables S8-S35 for details). The only unique donors are reported for the unassociated samples N1, N4 and N6.

|  | Reported donors |
| --- | --- |
| Outbreak and non-outbreak | *S.pseudointermedius* ED99 and HKU10-03<br>*S.warneri* SG1<br>*S.epidermidis* RP62A<br>*S.haemolyticus* JCSC1435<br>*S.aureus* COL<br>*S.lugdunensis* HKU09-01 |
| Non-outbreak only | *S.epidermidis* ATCC 12228 (N1,N6 only) and PM221 (N4 only)<br>*E.faecium* Aus0004 (N1 only) |

Table 4 states the total number of clustered HGT regions and the number of the clustered regions where HO 5096 0412 is the acceptor that are found by DaisySuite. Most HGT regions hence have the EMRSA-15 representative as acceptor.

Figure 4 shows the presence of the 41 HGT regions determined by mapping coverage called by Daisy among all samples. The purpose of the coverage analysis is to evaluate again if the HGT regions differ between the outbreak and non-outbreak strains but also to estimate if there are regions shared by all outbreak strains that are FN candidates of Daisy, or regions not covered at all that are likely FP candidates.

**Figure 4:**
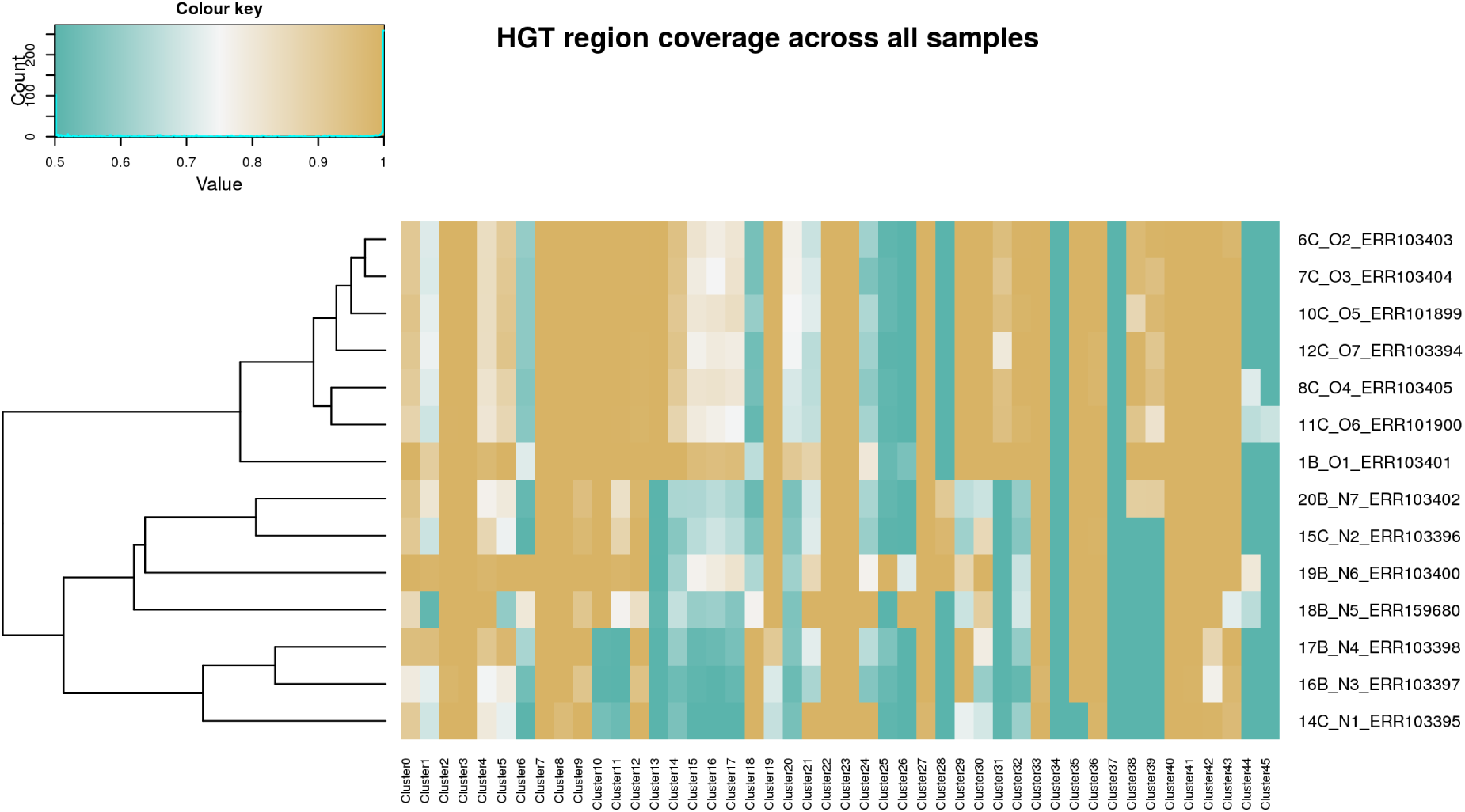
Heatmap of HGT region coverages. The mean coverages of HGT regions from all samples are calculated across every sample, and compared after application of a sigmoidal function. Solid green spots indicate no coverage, solid ochre high coverage. Regions 34 and 37 are not covered in any sample and hence FP calls. Sample O6 shows presence of multiple HGT regions called by DaisySuite for other samples but missed here. There is a distinct presence of HGT regions between the outbreak samples in the upper part and the unassociated samples in the lower part.

The clustering of samples according to the dendrogram shown in figure 4 was done automatically (see settings part), and hence reflects the relation of the samples according to the mapping coverage of the proposed HGT regions.

All outbreak strains are clustered together and share most of their HGT regions. All non-outbreak strains for which DaisyGPS did not report EMRSA-15 as an acceptor candidate are clustered away furthest from the outbreak strains (N1, N3-N5). The likely related samples N3 and N4 are clustered together. Regarding a distinction of outbreak and non-outbreak strains, DaisySuite is able to determine the outbreak-related HGT regions which differ from the HGT candidates for the non-outbreak strains. Hence, a distinction is possible. Although DaisySuite only called one HGT region for O6, we can deduce from the coverage profile that more HGT regions called for the other outbreak samples are present as well but were missed by DaisySuite. As can be seen in the heatmap, clusters 34 and 37 are not covered by any sample and hence likely FPs. We detected the AMR gene *mecA* on Cluster 0, however, resistance is shared among all 14 samples according to Köser et al. (2012). No further AMR genes tested by Köser et al. (2012) are detected on the other clusters. However, most of these AMR genes are on plasmids that were not analysed here.

## 4 Discussion

We presented DaisyGPS, a pipeline that facilitates metagenomic profiling strategies to identify acceptor and donor candidates from NGS reads of a potential HGT organism. DaisyGPS, together with Daisy, is part of the comprehensive HGT detection suite DaisySuite. We successfully validated DaisyGPS on simulated and real data previously analysed in Trappe et al. (2016). We further demonstrated robustness of the DaisySuite on a large-scale simulation with 100 negative HGT tests, showing that DaisySuite correctly reports no HGT events with a specificity of 97%. On a large-scale simulation with 100 positive HGT simulations, DaisySuite reports the correct HGT event with a total sensitivity of 69%. From the 818 pairs reported by DaisyGPS among the 100 simulations, Daisy called the TP and TN regions with a sensitivity of 89%. Lastly, we evaluated DaisySuite on an MRSA outbreak data set with seven outbreak associated samples and seven not associated with the outbreak but that occurred during the same time frame. Here we could show that DaisySuite successfully distinguishes between associated and not associated samples regarding their suggested HGT regions, i.e. the outbreak samples show a distinct number and content of reported HGT regions.

One has to acknowledge that all outbreak strains have a high sequence similarity to the EMRSA-15 strain, which is not necessarily the case for the non-outbreak strains. This is also reflected in the results from DaisyGPS where *S.aureus* HO 5096 0412 is the best acceptor candidate for all outbreak strains but not reported at all for some non-outbreak strains. It directly follows that a sequence comparison based analysis as done with DaisySuite will likely find different patterns for the outbreak and non-outbreak strains, and a difference in HGT region candidates might seem obvious. However, starting from having established such a difference, there is value in then analysing the shared HGT region candidates among the outbreak-related strains. For this proof of concept, we performed a relatively simple evaluation by performing a coverage analysis of all HGT regions across all samples and investigating the presence of AMR genes within the HGT regions. But a future thorough follow-up analysis of the origin and functionality provided by the potential HGT sites could benefit our understanding of the risk and pathogenicity of these outbreak strains.

The observed FP and FN candidates, however, also reveal weaknesses of the sequence comparison approach. DaisyGPS is designed with a focus on sensitivity and hence inevitably leads to FP acceptor and donor candidate pairs to be examined by Daisy. Since these FPs are still due to a sufficient degree of mapping coverage, spurious split-reads and spanning reads can cause down-stream FP calls as observed for the simulated data set from *E.coli K12* DH10 and *H.pylori*. The reported HGT site from *H.ducreyi* has only similarities in the start and end part of the proposed region compared to the transferred *H.pylori* region though. Insertion sites can also lie within repeat regions which enhances the negative impact of ambiguous mappings. This emphasises that a critical evaluation of HGT predictions is always crucial.

From the missing HGT region calls for sample O6 that could be inferred from the coverage analysis, we can deduce that DaisySuite does not detect all HGT regions due to insufficient evidence. A potential cause could be that DaisyGPS did not report the correct donor reference. Even if DaisyGPS could find an appropriate donor genome, it is still likely that the genome content differs between the region present in the donor and the region actually present in the HGT organism. An alternative, complementary approach to cope with this problem of a lack of a suitable donor candidate could be to facilitate local, insertion sequence assembly. By offering identified insertion sequences, we can still provide the content of a potential HGT sequence and thereby enable down-stream analysis. This approach would also support the detection of novel HGT sequences not present in current reference databases, and therefore also the detection of, e.g., novel antimicrobial resistance genes. Popins (Kehr et al., 2015) is a tool for population-based insertion calling developed for human sequencing data (see, e.g., Kehr et al. (2017)). Popins only locally assembles unmapped reads (same input as for Daisy) with Velvet guided by a reference, thereby minimising the risk of potential misassemblies. On top of the assembly, Popins first uses spanning pairs (see red read pairs in Figure 1) to place an insertion in the (acceptor) reference, and then performs a local split-read alignment around the potential breakpoint. If multiple samples are provided, Popins merges contigs across samples into supercontigs, assuming that the same insertion is present in multiple samples. Although different bacterial samples do not represent a population as given for human populations, outbreak related samples still resemble a population such that one could use Popins for this purpose and gain valuable information. However, local insertion assembly only gives evidence for an insertion compared to the chosen acceptor reference, that does not necessarily mean that the insertion resulted from an HGT event. Hence, means to sophistically include insertion assembly results into the HGT context need to be defined first. Despite the evidence for an HGT event that DaisySuite can provide, the results should always be tested for alternative causations such as gene loss.

## 5 Conclusion

With DaisyGPS, we present a tool for acceptor and donor identification from NGS reads of an HGT organism. To do that, DaisyGPS refines metrics already defined and used for metagenomic profiling purposes to account for the acceptor and donor specific coverage profiles. We integrated DaisyGPS with Daisy into a comprehensive HGT detection suite, called DaisySuite, that provides an automatic workflow to first determine acceptor and donor candidates and then identify and characterise HGT regions from the suggested acceptor-donor pairs. We successfully evaluated DaisyGPS on data previously analysed with Daisy, and demonstrated sensitivity and robustness of the DaisySuite in a largescale simulation with 100 simulated positive and negative HGT events. We could further show the benefits of an HGT analysis with DaisySuite on an MRSA outbreak data set where DaisySuite reported HGT candidates that help to distinguish between outbreak associated and unassociated samples and therefore also provide information for outbreak strain characterisation.

## Acknowledgement

We thank Tobias Marschall and Jan Rouven Forster for inspiring discussions.

## Author’s Contributions

KT, ES and BYR conceived the study and analysed data. KT and ES wrote the manuscript. ES developed and KT participated in developing the pipeline. BYR participated in manuscript editing. All authors read and approved the final manuscript.

---

**Table S1:** Confusion matrix for DaisyGPS classifications. If the simulation contains an HGT and DaisyGPS reports at least one candidate pair that corresponds to the correct acceptor/donor pair, the run is considered a TP. If DaisyGPS fails to report the correct acceptor or donor, the run is deemed a FP since all pairs will undergo follow up analysis by Daisy. In a negative test setting, a FP occurs if DaisyGPS reports any pair where the acceptor does not equal the donor and a TN means that either no pair was reported or acceptor and donor of the pair are the same organism.

|  |  | True condition (ground truth) |  |
| --- | --- | --- | --- |
|  |  | Simulation contains HGT<br>(positive setting) | Simulation does not contain HGT<br>(negative setting) |
| Predicted condition<br>(DaisyGPS) | Run reports HGT<br>Run does not report any HGT | TP<br>FP | FP<br>TN |

**Table S2:** Confusion matrix for DaisyGPS classifications. If the simulation contains an HGT and DaisyGPS reports at least one candidate pair that corresponds to the correct acceptor/donor pair, the run is considered a TP. If DaisyGPS fails to report the correct acceptor or donor, the run is deemed a FP since all pairs will undergo follow up analysis by Daisy. In a negative test setting, a FP occurs if DaisyGPS reports any pair where the acceptor does not equal the donor and a TN means that either no pair was reported or acceptor and donor of the pair are the same organism.

|  |  | True condition (ground truth) |  |
| --- | --- | --- | --- |
|  |  | Pair represents HGT<br>(DaisyGPS TP) | Pair does not represent HGT<br>(DaisyGPS FP) |
| Predicted condition<br>(Daisy) | Pair reports HGT<br>Pair does not report any HGT | TP<br>FP | FP<br>TN |

**Table S3:**
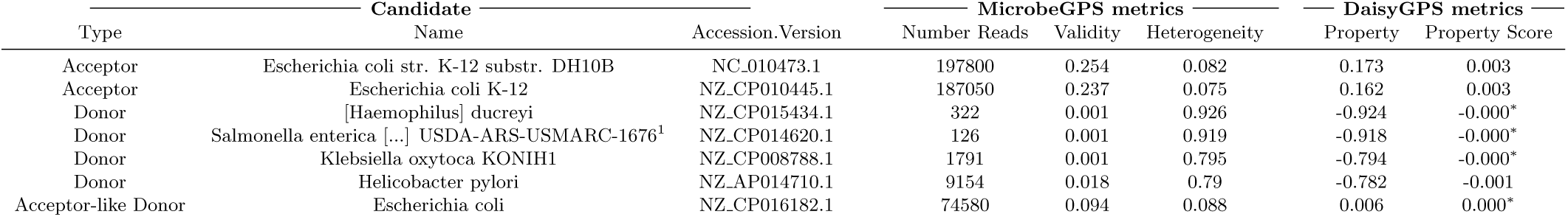
Acceptor and donor candidates for sim1HP run with yara, no species filter and no samflag filter. Sampling sensitivity = 90. No taxon blacklist. No parent blacklist. No species blacklist. (-)0.000 represents absolute values 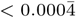.^1^Salmonella enterica subsp. enterica serovar Anatum str. USDA-ARS-USMARC-1676

**Table S4:** Results for sim1HP run with yara, gustaf, no species filter and no samflag filter. Sampling sensitivity = 90. Split read threshold = 3. No taxon blacklist. No parent blacklist. No species blacklist.

| Organism |  | Acceptor |  |  | Donor |  |  | Read Evidence |  |  | Evidence Filter |  |  |  |
| --- | --- | --- | --- | --- | --- | --- | --- | --- | --- | --- | --- | --- | --- | --- |
| Acceptor | Donor | Start | End | Coverage | Start | End | Coverage | Split | Spanning | Within | A-Cov | D-Cov | Spanning | Within |
| NZ_CP010445.1 | NZ_AP014710.1 | 1880235 | 1880237 | 44.0 | 1322002 | 1350000 | 94.62 | 152 | 182 | 8712 | 7 | 100 | 100 | 100 |
| NZ_CP010445.1 | NZ_CP015434.1 | 3904873 | 3904886 | 40.54 | 114928 | 126957 | 30.41 | 871 | 156 | 884 | 3 | 100 | 100 | 100 |
| NC_010473.1 | NZ_AP014710.1 | 1120261 | 1120263 | 43.0 | 1322002 | 1350000 | 94.62 | 154 | 182 | 8712 | 3 | 100 | 100 | 100 |

**Table S5:**
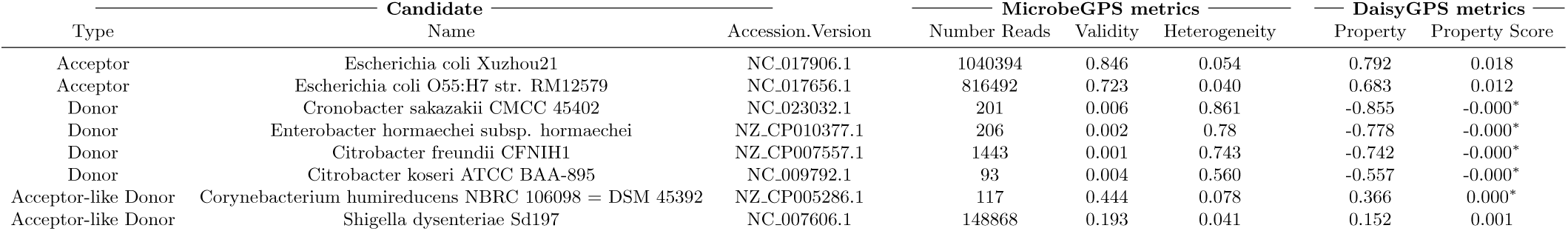
Acceptor and donor candidates for real1B run with yara, species filter and no samflag filter. Taxon blacklist: [83334, 1045010]. Parent blacklist: [83334]. No species blacklist. (-)0.000^*\**^represents absolute values 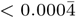.

**Table S6:** Results for real1B run with yara, gustaf, species filter and no samflag filter. Sampling sensitivity = 95. Split read threshold = 3. Taxon blacklist: [83334, 1045010]. Parent blacklist: [83334]. No species blacklist. Results (139 HGT candidates) for NC 017656.1 (acceptor) and NZ CP007557.1 (donor) are omitted here for sake of simplicity. For all other pairs no HGT candidates were reported.

| Organism |  | Acceptor |  |  | Donor |  |  | Read Evidence |  |  | Evidence Filter |  |  |  |
| --- | --- | --- | --- | --- | --- | --- | --- | --- | --- | --- | --- | --- | --- | --- |
| Acceptor | Donor | Start | End | Coverage | Start | End | Coverage | Split | Spanning | Within | A-Cov | D-Cov | Spanning | Within |
| NC_017656.1 | NC_007606.1 | 314439 | 334641 | 27.39 | 2213697 | 2214454 | 63.18 | 39 | 3 | 102 | 0 | 100 | 100 | 100 |
| NC_017656.1 | NC_007606.1 | 1570633 | 1580081 | 138.85 | 1282007 | 1320884 | 7.51 | 9 | 1 | 714 | 100 | 97 | 96 | 98 |
| NC_017656.1 | NC_007606.1 | 1570633 | 1584983 | 141.99 | 1282007 | 1329491 | 11.14 | 11 | 12 | 973 | 99 | 97 | 98 | 97 |
| NC_017656.1 | NC_007606.1 | 1580080 | 1584983 | 148.04 | 1320883 | 1329491 | 27.6 | 8 | 12 | 261 | 99 | 99 | 99 | 99 |
| NC_017656.1 | NC_007606.1 | 1589216 | 1618452 | 247.73 | 4032919 | 4035786 | 110.69 | 107 | 10 | 576 | 100 | 100 | 100 | 100 |
| NC_017656.1 | NC_007606.1 | 1738741 | 1739271 | 30.87 | 1321240 | 1322115 | 88.45 | 42 | 73 | 60 | 4 | 100 | 100 | 98 |
| NC_017656.1 | NC_007606.1 | 1738741 | 1739785 | 157.15 | 1321240 | 1322656 | 58.2 | 17 | 5 | 72 | 95 | 100 | 100 | 99 |
| NC_017656.1 | NC_007606.1 | 1738741 | 1740010 | 134.9 | 1321240 | 1322870 | 51.13 | 50 | 3 | 72 | 96 | 100 | 100 | 98 |
| NC_017656.1 | NC_007606.1 | 1738741 | 1740078 | 129.54 | 1321240 | 1322973 | 49.81 | 17 | 6 | 81 | 100 | 98 | 100 | 98 |
| NC_017656.1 | NC_007606.1 | 1738741 | 1745278 | 119.31 | 1321240 | 1331304 | 23.91 | 9 | 52 | 202 | 96 | 98 | 100 | 98 |
| NC_017656.1 | NC_007606.1 | 1739270 | 1739785 | 287.13 | 1322114 | 1322656 | 9.33 | 56 | 5 | 13 | 99 | 96 | 100 | 99 |
| NC_017656.1 | NC_007606.1 | 1739270 | 1740477 | 130.81 | 1322114 | 1323341 | 21.27 | 28 | 3 | 42 | 96 | 98 | 99 | 98 |
| NC_017656.1 | NC_007606.1 | 1739270 | 1745278 | 127.11 | 1322114 | 1331304 | 17.77 | 24 | 52 | 143 | 97 | 99 | 99 | 96 |
| NC_017656.1 | NC_007606.1 | 1739784 | 1741539 | 10.67 | 1283675 | 1322655 | 11.22 | 19 | 294 | 897 | 4 | 97 | 100 | 97 |
| NC_017656.1 | NC_007606.1 | 1739784 | 1745278 | 112.11 | 1322655 | 1331304 | 18.29 | 16 | 51 | 130 | 95 | 100 | 100 | 100 |
| NC_017656.1 | NC_007606.1 | 1740009 | 1740477 | 6.25 | 1322869 | 1323341 | 42.62 | 20 | 3 | 28 | 5 | 97 | 100 | 96 |
| NC_017656.1 | NC_007606.1 | 1740009 | 1745278 | 115.53 | 1322869 | 1331304 | 18.65 | 17 | 52 | 129 | 98 | 99 | 100 | 96 |
| NC_017656.1 | NC_007606.1 | 1740077 | 1740477 | 2.25 | 1322972 | 1323341 | 46.64 | 16 | 3 | 25 | 4 | 100 | 100 | 100 |
| NC_017656.1 | NC_007606.1 | **1741538** | **1744925** | 164.13 | **1283674** | **1288080** | 59.4 | 18 | 9 | 692 | 99 | 100 | 100 | 100 |
| NC_017656.1 | NC_007606.1 | 1741538 | 1745278 | 159.71 | 1283674 | 1331304 | 12.51 | 9 | 166 | 1031 | 100 | 97 | 99 | 95 |
| NC_017656.1 | NC_007606.1 | 1957909 | 1958879 | 132.94 | 4032919 | 4035786 | 110.69 | 41 | 7 | 576 | 99 | 99 | 98 | 99 |
| NC_017656.1 | NC_007606.1 | 1957909 | 1982375 | 118.01 | 4032919 | 4035782 | 110.56 | 17 | 12 | 576 | 97 | 100 | 100 | 100 |
| NC_017656.1 | NC_007606.1 | 1958870 | 1982375 | 117.37 | 4034933 | 4035782 | 356.29 | 22 | 35 | 576 | 98 | 100 | 100 | 100 |
| NC_017656.1 | NC_007606.1 | 1986050 | 1986053 | 726.33 | 1288361 | 1331322 | 7.47 | 10 | 335 | 319 | 100 | 97 | 100 | 95 |
| NC_017656.1 | NC_007606.1 | 1986050 | 1992463 | 155.63 | 1321775 | 1331322 | 25.25 | 126 | 72 | 197 | 99 | 98 | 100 | 97 |
| NC_017656.1 | NC_007606.1 | 1986234 | 1992463 | 146.03 | 1321775 | 1329808 | 28.06 | 261 | 80 | 190 | 100 | 98 | 100 | 96 |
| NC_017656.1 | NC_007606.1 | 1986234 | 1992955 | 155.68 | 1320887 | 1329808 | 32.55 | 35 | 126 | 308 | 99 | 100 | 100 | 99 |
| NC_017656.1 | NC_007606.1 | 1992462 | 1992955 | 277.57 | 1320887 | 1321774 | 73.17 | 131 | 91 | 106 | 100 | 99 | 100 | 99 |
| NC_017656.1 | NC_007606.1 | 2431977 | 2443616 | 15.53 | 1282008 | 1322832 | 10.76 | 17 | 60 | 897 | 0 | 96 | 100 | 96 |
| NC_017656.1 | NC_007606.1 | 2435781 | 2443492 | 8.8 | 1282069 | 1320883 | 7.51 | 193 | 62 | 714 | 3 | 98 | 98 | 98 |
| NC_017656.1 | NC_007606.1 | 2469232 | 2481815 | 49.5 | 4032919 | 4035785 | 110.66 | 81 | 24 | 576 | 2 | 99 | 100 | 99 |
| NC_017656.1 | NC_007606.1 | 2486033 | 2488461 | 149.98 | 4298967 | 4301718 | 16.95 | 23 | 5 | 67 | 95 | 97 | 100 | 96 |
| NC_017656.1 | NC_007606.1 | 2486033 | 2488662 | 150.6 | 4298967 | 4301905 | 16.19 | 65 | 10 | 68 | 99 | 98 | 100 | 98 |
| NC_017656.1 | NC_007606.1 | 2486203 | 2488662 | 153.24 | 4299043 | 4301905 | 16.62 | 47 | 10 | 68 | 99 | 97 | 100 | 95 |
| NC_017656.1 | NC_007606.1 | 2486203 | 2488723 | 152.86 | 4299043 | 4301977 | 17.39 | 29 | 10 | 69 | 98 | 96 | 100 | 95 |
| NC_017656.1 | NC_007606.1 | 2487505 | 2489413 | 150.49 | 953376 | 956244 | 23.61 | 10 | 3 | 119 | 98 | 98 | 99 | 99 |
| NC_017656.1 | NC_007606.1 | 2488461 | 2489413 | 130.37 | 953376 | 954653 | 52.39 | 12 | 4 | 119 | 95 | 99 | 100 | 97 |
| NC_017656.1 | NC_007606.1 | 2488601 | 2488723 | 136.13 | 4301842 | 4301977 | 32.39 | 8 | 11 | 2 | 98 | 97 | 100 | 96 |
| NC_017656.1 | NC_007606.1 | 2678766 | 2679015 | 44.61 | 1323123 | 1323370 | 29.6 | 18 | 4 | 5 | 5 | 97 | 100 | 96 |
| NC_017656.1 | NC_007606.1 | 3607310 | 3629241 | 31.35 | 4189699 | 4189800 | 491.86 | 42 | 24 | 12 | 0 | 100 | 100 | 98 |
| NC_017656.1 | NC_007606.1 | 3615738 | 3630353 | 153.33 | 4195901 | 4198011 | 700.99 | 149 | 6 | 4245 | 97 | 100 | 98 | 100 |
| NC_017656.1 | NC_007606.1 | 3615738 | 3632904 | 131.04 | 4195901 | 4206697 | 139.78 | 21 | 4 | 4245 | 96 | 100 | 98 | 100 |
| NC_017656.1 | NC_007606.1 | 3615738 | 3632993 | 130.65 | 4195901 | 4206818 | 138.38 | 19 | 4 | 4250 | 98 | 100 | 98 | 100 |
| NC_017656.1 | NC_007606.1 | 3629240 | 3630353 | 1409.38 | 4189698 | 4198011 | 184.22 | 222 | 38 | 4278 | 100 | 100 | 100 | 100 |
| NC_017656.1 | NC_007606.1 | 3629240 | 3632904 | 430.46 | 4189698 | 4206697 | 91.85 | 30 | 36 | 4278 | 100 | 100 | 100 | 100 |
| NC_017656.1 | NC_007606.1 | 3629240 | 3632993 | 421.57 | 4189698 | 4206818 | 91.3 | 27 | 36 | 4283 | 100 | 100 | 100 | 100 |

**Table S7:** Acceptor and donor candidates for real4 run with yara, no species filter and no samflag filter. Taxon blacklist: [595495]. No parent blacklist. No species blacklist. (-)0.000^*\**^represents absolute values 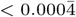.

| Type | Candidate |  | MicrobeGPS metrics |  |  | DaisyGPS metrics |  |
| --- | --- | --- | --- | --- | --- | --- | --- |
|  | Name | Accession.Version | Number Reads | Validity | Heterogeneity | Property | Property Score |
| Acceptor | Escherichia coli W | NC_017635.1 | 221389 | 0.852 | 0.024 | 0.829 | 0.026 |
| Acceptor | Escherichia coli W | NC_017664.1 | 221570 | 0.853 | 0.025 | 0.828 | 0.026 |
| Donor | Salmonella enterica subsp. enterica serovar Infantis | NZ_CP016410.1 | 83 | 0.005 | 0.943 | -0.938 | -0.000* |
| Donor | [Haemophilus] ducreyi | NZ_CP015434.1 | 119 | 0.001 | 0.920 | -0.919 | -0.000* |
| Donor | Zymomonas mobilis subsp. mobilis NRRL B-12526 | NZ_CP003709.1 | 3067 | 0.002 | 0.876 | -0.874 | -0.000* |
| Acceptor-like Donor | Shigella boydii CDC 3083-94 | NC_010658.1 | 23506 | 0.150 | 0.047 | 0.104 | 0.000* |
| Acceptor-like Donor | Shigella sonnei 53G | NC_016822.1 | 29127 | 0.168 | 0.073 | 0.095 | 0.000* |

**Table S8:** Acceptor and donor candidates for ERR103401 run with yara, species filter and no samflag filter. No taxon blacklist. No parent blacklist. No species blacklist. (-)0.000^*\**^represents absolute values 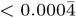.

| Type | Candidate |  | MicrobeGPS metrics |  |  | DaisyGPS metrics |  |
| --- | --- | --- | --- | --- | --- | --- | --- |
|  | Name | Accession.Version | Number Reads | Validity | Heterogeneity | Property | Property Score |
| Acceptor | Staphylococcus aureus subsp. aureus HO 5096 0412 | NC_017763.1 | 440076 | 0.832 | 0.04 | 0.792 | 0.041 |
| Acceptor | Staphylococcus aureus subsp. aureus | NZ_CP007659.1 | 439586 | 0.824 | 0.041 | 0.783 | 0.040 |
| Donor | Staphylococcus pseudintermedius ED99 | NC_017568.1 | 1089 | 0.002 | 0.691 | -0.689 | -0.000* |
| Donor | Staphylococcus warneri SG1 | NC_020164.1 | 523 | 0.003 | 0.631 | -0.628 | -0.000* |
| Donor | Staphylococcus epidermidis RP62A | NC_002976.3 | 5512 | 0.006 | 0.540 | -0.534 | -0.000* |
| Donor | Staphylococcus haemolyticus JCSC1435 | NC_007168.1 | 3614 | 0.005 | 0.291 | -0.285 | -0.000* |
| Donor | Staphylococcus aureus subsp. aureus COL | NC_002951.2 | 49889 | 0.106 | 0.233 | -0.127 | -0.001 |
| Acceptor-like Donor | Staphylococcus aureus subsp. aureus | NZ_CP012011.1 | 54992 | 0.11 | 0.109 | 0.001 | 0.000* |

**Table S9:**
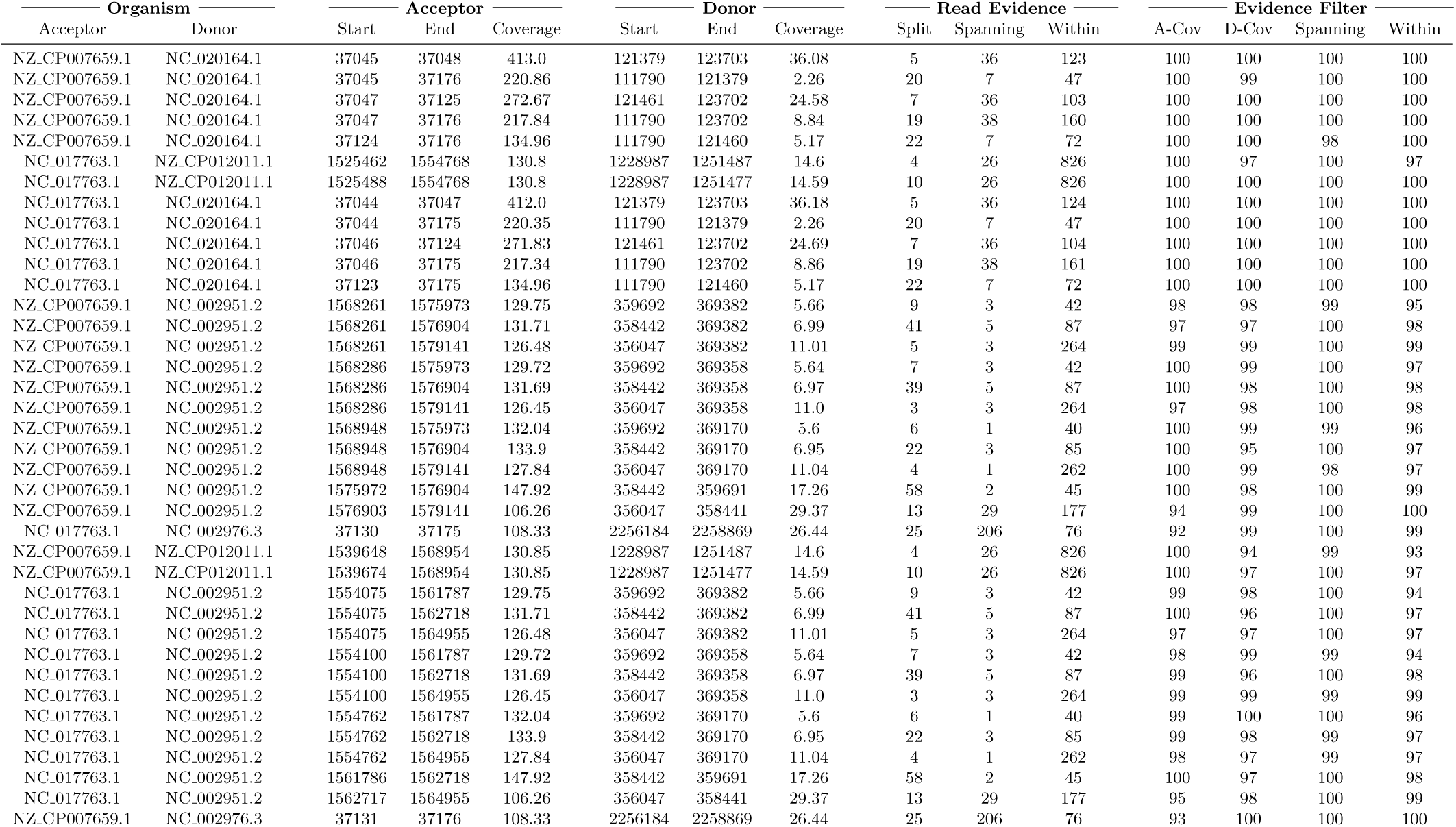
Results for ERR103401 run with yara, gustaf, species filter and no samflag filter. Sampling sensitivity = 90. Split read threshold = 3. No taxon blacklist. No parent blacklist. No species blacklist.

**Table S10:**
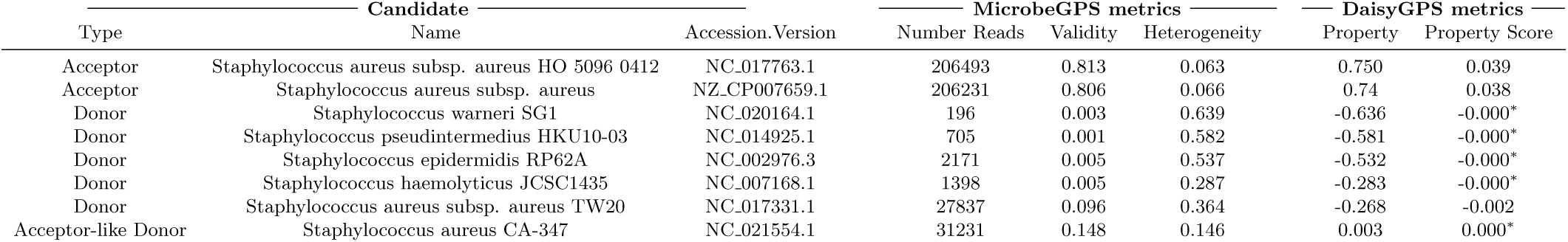
Acceptor and donor candidates for ERR103403 run with yara, species filter and no samflag filter. Sampling sensitivity = 85. No taxon blacklist. No parent blacklist. No species blacklist. (-)0.000^*\**^represents absolute values 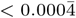.

**Table S11:**
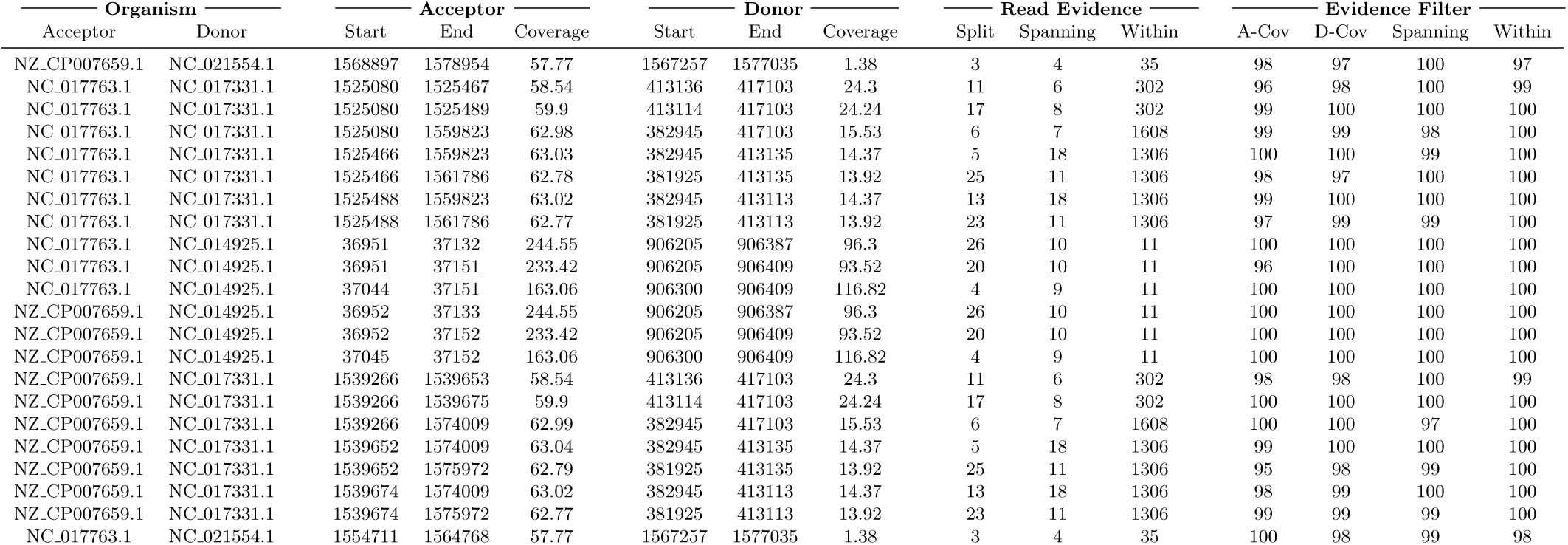
Results for ERR103403 run with yara, gustaf, species filter and no samflag filter. Sampling sensitivity = 90. Split read threshold = 3. No taxon blacklist. No parent blacklist. No species blacklist.

| Organism |  | Acceptor |  |  | Donor |  |  | Read Evidence |  |  | Evidence Filter |  |  |  |
| --- | --- | --- | --- | --- | --- | --- | --- | --- | --- | --- | --- | --- | --- | --- |
| Acceptor | Donor | Start | End | Coverage | Start | End | Coverage | Split | Spanning | Within | A-Cov | D-Cov | Spanning | Within |
| NZ_CP007659.1 | NC_021554.1 | 1568897 | 1578954 | 57.77 | 1567257 | 1577035 | 1.38 | 3 | 4 | 35 | 98 | 97 | 100 | 97 |
| NC_017763.1 | NC_017331.1 | 1525080 | 1525467 | 58.54 | 413136 | 417103 | 24.3 | 11 | 6 | 302 | 96 | 98 | 100 | 99 |
| NC_017763.1 | NC_017331.1 | 1525080 | 1525489 | 59.9 | 413114 | 417103 | 24.24 | 17 | 8 | 302 | 99 | 100 | 100 | 100 |
| NC_017763.1 | NC_017331.1 | 1525080 | 1559823 | 62.98 | 382945 | 417103 | 15.53 | 6 | 7 | 1608 | 99 | 99 | 98 | 100 |
| NC_017763.1 | NC_017331.1 | 1525466 | 1559823 | 63.03 | 382945 | 413135 | 14.37 | 5 | 18 | 1306 | 100 | 100 | 99 | 100 |
| NC_017763.1 | NC_017331.1 | 1525466 | 1561786 | 62.78 | 381925 | 413135 | 13.92 | 25 | 11 | 1306 | 98 | 97 | 100 | 100 |
| NC_017763.1 | NC_017331.1 | 1525488 | 1559823 | 63.02 | 382945 | 413113 | 14.37 | 13 | 18 | 1306 | 99 | 100 | 100 | 100 |
| NC_017763.1 | NC_017331.1 | 1525488 | 1561786 | 62.77 | 381925 | 413113 | 13.92 | 23 | 11 | 1306 | 97 | 99 | 99 | 100 |
| NC_017763.1 | NC_014925.1 | 36951 | 37132 | 244.55 | 906205 | 906387 | 96.3 | 26 | 10 | 11 | 100 | 100 | 100 | 100 |
| NC_017763.1 | NC_014925.1 | 36951 | 37151 | 233.42 | 906205 | 906409 | 93.52 | 20 | 10 | 11 | 96 | 100 | 100 | 100 |
| NC_017763.1 | NC_014925.1 | 37044 | 37151 | 163.06 | 906300 | 906409 | 116.82 | 4 | 9 | 11 | 100 | 100 | 100 | 100 |
| NZ_CP007659.1 | NC_014925.1 | 36952 | 37133 | 244.55 | 906205 | 906387 | 96.3 | 26 | 10 | 11 | 100 | 100 | 100 | 100 |
| NZ_CP007659.1 | NC_014925.1 | 36952 | 37152 | 233.42 | 906205 | 906409 | 93.52 | 20 | 10 | 11 | 100 | 100 | 100 | 100 |
| NZ_CP007659.1 | NC_014925.1 | 37045 | 37152 | 163.06 | 906300 | 906409 | 116.82 | 4 | 9 | 11 | 100 | 100 | 100 | 100 |
| NZ_CP007659.1 | NC_017331.1 | 1539266 | 1539653 | 58.54 | 413136 | 417103 | 24.3 | 11 | 6 | 302 | 98 | 98 | 100 | 99 |
| NZ_CP007659.1 | NC_017331.1 | 1539266 | 1539675 | 59.9 | 413114 | 417103 | 24.24 | 17 | 8 | 302 | 100 | 100 | 100 | 100 |
| NZ_CP007659.1 | NC_017331.1 | 1539266 | 1574009 | 62.99 | 382945 | 417103 | 15.53 | 6 | 7 | 1608 | 100 | 100 | 97 | 100 |
| NZ_CP007659.1 | NC_017331.1 | 1539652 | 1574009 | 63.04 | 382945 | 413135 | 14.37 | 5 | 18 | 1306 | 99 | 100 | 100 | 100 |
| NZ_CP007659.1 | NC_017331.1 | 1539652 | 1575972 | 62.79 | 381925 | 413135 | 13.92 | 25 | 11 | 1306 | 95 | 98 | 99 | 100 |
| NZ_CP007659.1 | NC_017331.1 | 1539674 | 1574009 | 63.02 | 382945 | 413113 | 14.37 | 13 | 18 | 1306 | 98 | 99 | 100 | 100 |
| NZ_CP007659.1 | NC_017331.1 | 1539674 | 1575972 | 62.77 | 381925 | 413113 | 13.92 | 23 | 11 | 1306 | 99 | 99 | 99 | 100 |
| NC_017763.1 | NC_021554.1 | 1554711 | 1564768 | 57.77 | 1567257 | 1577035 | 1.38 | 3 | 4 | 35 | 100 | 98 | 99 | 98 |

**Table S12:** Acceptor and donor candidates for ERR103404 run with yara, species filter and no samflag filter. Sampling sensitivity = 85. No taxon blacklist. No parent blacklist. No species blacklist. (-)0.000^*\**^represents absolute values 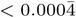.

| Candidate |  |  | MicrobeGPS metrics |  |  | DaisyGPS metrics |  |
| --- | --- | --- | --- | --- | --- | --- | --- |
| Type | Name | Accession.Version | Number Reads | Validity | Heterogeneity | Property | Property Score |
| Acceptor | Staphylococcus aureus subsp. aureus HO 5096 0412 | NC_017763.1 | 193345 | 0.812 | 0.043 | 0.769 | 0.041 |
| Acceptor | Staphylococcus aureus subsp. aureus | NZ_CP007659.1 | 193065 | 0.805 | 0.044 | 0.761 | 0.041 |
| Donor | Staphylococcus pseudintermedius ED99 | NC_017568.1 | 459 | 0.001 | 0.702 | -0.700 | -0.000* |
| Donor | Staphylococcus warneri SG1 | NC_020164.1 | 244 | 0.003 | 0.631 | -0.627 | -0.000* |
| Donor | Staphylococcus epidermidis RP62A | NC_002976.3 | 2256 | 0.005 | 0.536 | -0.531 | -0.000* |
| Donor | Staphylococcus haemolyticus JCSC1435 | NC_007168.1 | 1441 | 0.005 | 0.299 | -0.295 | -0.000* |
| Donor | Staphylococcus aureus subsp. aureus COL | NC_002951.2 | 20891 | 0.101 | 0.233 | -0.133 | -0.001 |
| Acceptor-like Donor | Staphylococcus aureus subsp. aureus DSM 20231 | NZ_CP011526.1 | 16400 | 0.102 | 0.084 | 0.018 | 0.000* |

**Table S13:**
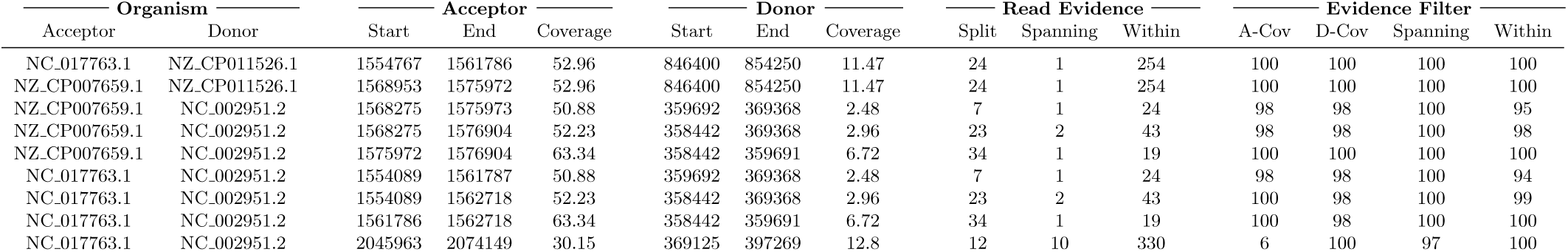
Results for ERR103404 run with yara, gustaf, species filter and no samflag filter. Sampling sensitivity = 90. Split read threshold = 3. No taxon blacklist. No parent blacklist. No species blacklist.

| Organism |  | Acceptor |  |  | Donor |  |  | Read Evidence |  |  | Evidence Filter |  |  |  |
| --- | --- | --- | --- | --- | --- | --- | --- | --- | --- | --- | --- | --- | --- | --- |
| Acceptor | Donor | Start | End | Coverage | Start | End | Coverage | Split | Spanning | Within | A-Cov | D-Cov | Spanning | Within |
| NC_017763.1 | NZ_CP011526.1 | 1554767 | 1561786 | 52.96 | 846400 | 854250 | 11.47 | 24 | 1 | 254 | 100 | 100 | 100 | 100 |
| NZ_CP007659.1 | NZ_CP011526.1 | 1568953 | 1575972 | 52.96 | 846400 | 854250 | 11.47 | 24 | 1 | 254 | 100 | 100 | 100 | 100 |
| NZ_CP007659.1 | NC_002951.2 | 1568275 | 1575973 | 50.88 | 359692 | 369368 | 2.48 | 7 | 1 | 24 | 98 | 98 | 100 | 95 |
| NZ_CP007659.1 | NC_002951.2 | 1568275 | 1576904 | 52.23 | 358442 | 369368 | 2.96 | 23 | 2 | 43 | 98 | 98 | 100 | 98 |
| NZ_CP007659.1 | NC_002951.2 | 1575972 | 1576904 | 63.34 | 358442 | 359691 | 6.72 | 34 | 1 | 19 | 100 | 100 | 100 | 100 |
| NC_017763.1 | NC_002951.2 | 1554089 | 1561787 | 50.88 | 359692 | 369368 | 2.48 | 7 | 1 | 24 | 98 | 98 | 100 | 94 |
| NC_017763.1 | NC_002951.2 | 1554089 | 1562718 | 52.23 | 358442 | 369368 | 2.96 | 23 | 2 | 43 | 100 | 98 | 100 | 99 |
| NC_017763.1 | NC_002951.2 | 1561786 | 1562718 | 63.34 | 358442 | 359691 | 6.72 | 34 | 1 | 19 | 100 | 98 | 100 | 100 |
| NC_017763.1 | NC_002951.2 | 2045963 | 2074149 | 30.15 | 369125 | 397269 | 12.8 | 12 | 10 | 330 | 6 | 100 | 97 | 100 |

**Table S14:** Acceptor and donor candidates for ERR103405 run with yara, species filter and no samflag filter. Sampling sensitivity = 85. No taxon blacklist. No parent blacklist. No species blacklist. (-)0.000^*\**^represents absolute values 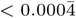.

| Candidate |  |  | MicrobeGPS metrics |  |  | DaisyGPS metrics |  |
| --- | --- | --- | --- | --- | --- | --- | --- |
| Type | Name | Accession.Version | Number Reads | Validity | Heterogeneity | Property | Property Score |
| Acceptor | Staphylococcus aureus subsp. aureus HO 5096 0412 | NC_017763.1 | 192851 | 0.811 | 0.03 | 0.781 | 0.041 |
| Acceptor | Staphylococcus aureus subsp. aureus | NZ_CP007659.1 | 192626 | 0.804 | 0.031 | 0.773 | 0.040 |
| Donor | Staphylococcus pseudintermedius ED99 | NC_017568.1 | 459 | 0.001 | 0.698 | -0.696 | -0.000* |
| Donor | Staphylococcus warneri SG1 | NC_020164.1 | 236 | 0.003 | 0.658 | -0.655 | -0.000* |
| Donor | Staphylococcus epidermidis RP62A | NC_002976.3 | 2006 | 0.005 | 0.543 | -0.538 | -0.000* |
| Donor | Staphylococcus haemolyticus JCSC1435 | NC_007168.1 | 1278 | 0.005 | 0.293 | -0.289 | -0.000* |
| Donor | Staphylococcus aureus subsp. aureus COL | NC_002951.2 | 21599 | 0.097 | 0.227 | -0.13 | -0.001 |
| Acceptor-like Donor | Staphylococcus aureus subsp. aureus | NZ_CP018205.1 | 20618 | 0.100 | 0.091 | 0.009 | 0.000* |

**Table S15:** Results for ERR103405 run with yara, gustaf, species filter and no samflag filter. Sampling sensitivity = 90. Split read threshold = 3. No taxon blacklist. No parent blacklist. No species blacklist.

| Organism |  | Acceptor |  |  | Donor |  |  | Read Evidence |  |  | Evidence Filter |  |  |  |
| --- | --- | --- | --- | --- | --- | --- | --- | --- | --- | --- | --- | --- | --- | --- |
| Acceptor | Donor | Start | End | Coverage | Start | End | Coverage | Split | Spanning | Within | A-Cov | D-Cov | Spanning | Within |
| NC_017763.1 | NZ_CP018205.1 | 1559883 | 1562718 | 58.73 | 1959491 | 1961823 | 7.95 | 12 | 1 | 38 | 100 | 100 | 100 | 100 |
| NC_017763.1 | NZ_CP018205.1 | 1561784 | 1562718 | 66.49 | 1960572 | 1961823 | 9.45 | 66 | 1 | 31 | 100 | 100 | 100 | 100 |
| NZ_CP007659.1 | NZ_CP018205.1 | 1574069 | 1576904 | 58.73 | 1959491 | 1961823 | 7.95 | 12 | 1 | 38 | 99 | 99 | 100 | 98 |
| NZ_CP007659.1 | NZ_CP018205.1 | 1575970 | 1576904 | 66.49 | 1960572 | 1961823 | 9.45 | 66 | 1 | 31 | 100 | 100 | 100 | 100 |
| NZ_CP007659.1 | NC_002951.2 | 1568261 | 1576904 | 56.11 | 358442 | 369382 | 3.13 | 10 | 1 | 50 | 100 | 98 | 100 | 100 |
| NZ_CP007659.1 | NC_002951.2 | 1575976 | 1576904 | 66.66 | 358442 | 359692 | 9.25 | 19 | 1 | 29 | 100 | 99 | 100 | 99 |
| NZ_CP007659.1 | NC_002951.2 | 2059982 | 2087935 | 30.68 | 369359 | 397269 | 12.87 | 5 | 12 | 341 | 10 | 100 | 100 | 100 |
| NZ_CP007659.1 | NC_002951.2 | 2059982 | 2088169 | 30.68 | 369125 | 397269 | 12.79 | 21 | 12 | 341 | 6 | 100 | 98 | 100 |
| NC_017763.1 | NC_002951.2 | 1554075 | 1562718 | 56.11 | 358442 | 369382 | 3.13 | 10 | 1 | 50 | 100 | 99 | 100 | 100 |
| NC_017763.1 | NC_002951.2 | 1561790 | 1562718 | 66.66 | 358442 | 359692 | 9.25 | 19 | 1 | 29 | 100 | 100 | 100 | 100 |
| NC_017763.1 | NC_002951.2 | 2045963 | 2073915 | 30.7 | 369359 | 397269 | 13.02 | 5 | 12 | 355 | 6 | 100 | 99 | 100 |
| NC_017763.1 | NC_002951.2 | 2045963 | 2074149 | 30.7 | 369125 | 397269 | 12.94 | 21 | 12 | 355 | 8 | 100 | 97 | 100 |

**Table S16:** Acceptor and donor candidates for ERR101899 run with yara, species filter and no samflag filter. Sampling sensitivity = 85. No taxon blacklist. No parent blacklist. No species blacklist. (-)0.000^*\**^represents absolute values 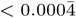.

| Candidate |  |  | MicrobeGPS metrics |  |  | DaisyGPS metrics |  |
| --- | --- | --- | --- | --- | --- | --- | --- |
| Type | Name | Accession.Version | Number Reads | Validity | Heterogeneity | Property | Property Score |
| Acceptor | Staphylococcus aureus subsp. aureus HO 5096 0412 | NC_017763.1 | 206272 | 0.814 | 0.047 | 0.767 | 0.040 |
| Acceptor | Staphylococcus aureus subsp. aureus | NZ_CP007659.1 | 206076 | 0.807 | 0.049 | 0.759 | 0.04 |
| Donor | Staphylococcus pseudintermedius ED99 | NC_017568.1 | 536 | 0.001 | 0.707 | -0.705 | -0.000* |
| Donor | Staphylococcus warneri SG1 | NC_020164.1 | 263 | 0.003 | 0.658 | -0.655 | -0.000* |
| Donor | Staphylococcus epidermidis RP62A | NC_002976.3 | 2226 | 0.005 | 0.537 | -0.532 | -0.000* |
| Donor | Staphylococcus haemolyticus JCSC1435 | NC_007168.1 | 1378 | 0.004 | 0.296 | -0.291 | -0.000* |
| Donor | Staphylococcus aureus subsp. aureus COL | NC_002951.2 | 22973 | 0.098 | 0.236 | -0.139 | -0.001 |
| Acceptor-like Donor | Staphylococcus aureus subsp. aureus DSM 20231 | NZ_CP011526.1 | 18223 | 0.099 | 0.085 | 0.014 | 0.000* |

**Table S17:** Results for ERR101899 run with yara, gustaf, species filter and no samflag filter. Sampling sensitivity = 90. Split read threshold = 3. No taxon blacklist. No parent blacklist. No species blacklist.

| Organism |  | Acceptor |  |  | Donor |  |  | Read Evidence |  |  | Evidence Filter |  |  |  |
| --- | --- | --- | --- | --- | --- | --- | --- | --- | --- | --- | --- | --- | --- | --- |
| Acceptor | Donor | Start | End | Coverage | Start | End | Coverage | Split | Spanning | Within | A-Cov | D-Cov | Spanning | Within |
| NZ_CP007659.1 | NC_002951.2 | 1568261 | 1575972 | 53.51 | 359694 | 369382 | 2.62 | 3 | 2 | 15 | 99 | 100 | 100 | 97 |
| NZ_CP007659.1 | NC_002951.2 | 1568261 | 1576904 | 55.07 | 358442 | 369382 | 3.05 | 8 | 2 | 34 | 99 | 99 | 99 | 99 |
| NZ_CP007659.1 | NC_002951.2 | 1568287 | 1575972 | 53.49 | 359694 | 369357 | 2.61 | 3 | 2 | 15 | 98 | 100 | 99 | 99 |
| NZ_CP007659.1 | NC_002951.2 | 1568287 | 1576904 | 55.06 | 358442 | 369357 | 3.04 | 8 | 2 | 34 | 100 | 100 | 99 | 99 |
| NZ_CP007659.1 | NC_002951.2 | 2059982 | 2087936 | 31.07 | 369358 | 397269 | 13.23 | 9 | 15 | 395 | 10 | 100 | 100 | 100 |
| NZ_CP007659.1 | NC_002951.2 | 2059982 | 2088169 | 31.08 | 369125 | 397269 | 13.15 | 31 | 15 | 396 | 7 | 100 | 94 | 100 |
| NZ_CP007659.1 | NC_020164.1 | 37045 | 37177 | 87.31 | 111789 | 121379 | 0.85 | 4 | 2 | 20 | 100 | 97 | 100 | 100 |
| NZ_CP007659.1 | NZ_CP011526.1 | 1568903 | 1575972 | 56.42 | 846397 | 854374 | 12.62 | 9 | 1 | 267 | 100 | 100 | 100 | 100 |
| NC_017763.1 | NZ_CP011526.1 | 1554717 | 1561786 | 56.42 | 846397 | 854374 | 12.62 | 9 | 1 | 267 | 98 | 100 | 100 | 100 |
| NC_017763.1 | NC_002951.2 | 1554075 | 1561786 | 53.51 | 359694 | 369382 | 2.62 | 3 | 2 | 15 | 97 | 98 | 100 | 94 |
| NC_017763.1 | NC_002951.2 | 1554075 | 1562718 | 55.07 | 358442 | 369382 | 3.05 | 8 | 2 | 34 | 99 | 99 | 100 | 100 |
| NC_017763.1 | NC_002951.2 | 1554101 | 1561786 | 53.49 | 359694 | 369357 | 2.61 | 3 | 2 | 15 | 98 | 99 | 99 | 93 |
| NC_017763.1 | NC_002951.2 | 1554101 | 1562718 | 55.06 | 358442 | 369357 | 3.04 | 8 | 2 | 34 | 99 | 98 | 100 | 97 |
| NC_017763.1 | NC_002951.2 | 2045963 | 2073916 | 31.1 | 369358 | 397269 | 13.45 | 9 | 15 | 415 | 8 | 100 | 91 | 100 |
| NC_017763.1 | NC_020164.1 | 37044 | 37176 | 87.31 | 111789 | 121379 | 0.85 | 4 | 2 | 20 | 100 | 100 | 99 | 100 |

**Table S18:**
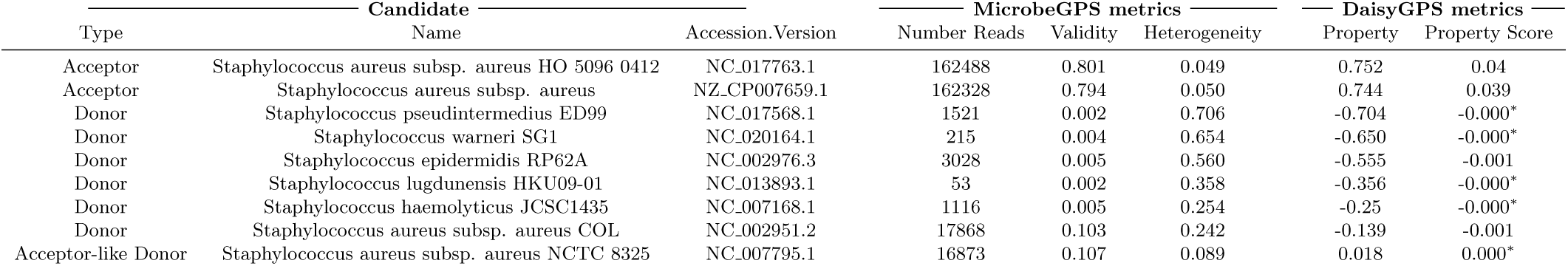
Acceptor and donor candidates for ERR101900 run with yara, species filter and no samflag filter. Sampling sensitivity = 85. No taxon blacklist. No parent blacklist. No species blacklist. (-)0.000^*\**^represents absolute values 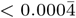.

**Table S19:** Results for ERR101900 run with yara, gustaf, species filter and no samflag filter. Sampling sensitivity = 90. Split read threshold = 3. No taxon blacklist. No parent blacklist. No species blacklist.

| Organism |  | Acceptor |  |  | Donor |  |  | Read Evidence |  |  | Evidence Filter |  |  |  |
| --- | --- | --- | --- | --- | --- | --- | --- | --- | --- | --- | --- | --- | --- | --- |
| Acceptor | Donor | Start | End | Coverage | Start | End | Coverage | Split | Spanning | Within | A-Cov | D-Cov | Spanning | Within |
| NC_017763.1 | NC_002951.2 | 1554089 | 1562718 | 45.31 | 358442 | 369368 | 2.81 | 15 | 1 | 31 | 100 | 100 | 99 | 98 |
| NC_017763.1 | NC_002951.2 | 1554762 | 1562718 | 46.77 | 358442 | 369170 | 2.78 | 8 | 1 | 29 | 100 | 100 | 98 | 98 |
| NC_017763.1 | NC_002951.2 | 1561790 | 1562718 | 53.62 | 358442 | 359696 | 5.56 | 8 | 1 | 15 | 98 | 99 | 100 | 100 |
| NZ_CP007659.1 | NC_007795.1 | 1575971 | 1576904 | 53.59 | 1961777 | 1963027 | 6.06 | 57 | 1 | 16 | 99 | 99 | 100 | 99 |
| NC_017763.1 | NC_007795.1 | 1561785 | 1562718 | 53.59 | 1961777 | 1963027 | 6.06 | 57 | 1 | 16 | 99 | 97 | 99 | 97 |
| NZ_CP007659.1 | NC_002951.2 | 1568275 | 1576904 | 45.31 | 358442 | 369368 | 2.81 | 15 | 1 | 31 | 99 | 99 | 100 | 99 |
| NZ_CP007659.1 | NC_002951.2 | 1568948 | 1576904 | 46.77 | 358442 | 369170 | 2.78 | 8 | 1 | 29 | 99 | 99 | 100 | 99 |
| NZ_CP007659.1 | NC_002951.2 | 1575976 | 1576904 | 53.62 | 358442 | 359696 | 5.56 | 8 | 1 | 15 | 100 | 99 | 100 | 98 |

**Table S20:**
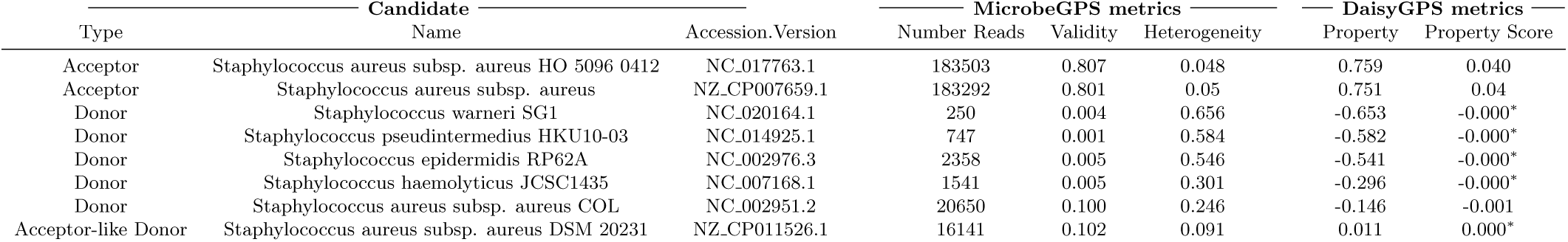
Acceptor and donor candidates for ERR103394 run with yara, species filter and no samflag filter. Sampling sensitivity = 85. No taxon blacklist. No parent blacklist. No species blacklist. (-)0.000^*\**^represents absolute values 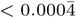.

**Table S21:** Results for ERR103394 run with yara, gustaf, species filter and no samflag filter. Sampling sensitivity = 90. Split read threshold = 3. No taxon blacklist. No parent blacklist. No species blacklist.

| Organism |  | Acceptor |  |  | Donor |  |  | Read Evidence |  |  | Evidence Filter |  |  |  |
| --- | --- | --- | --- | --- | --- | --- | --- | --- | --- | --- | --- | --- | --- | --- |
| Acceptor | Donor | Start | End | Coverage | Start | End | Coverage | Split | Spanning | Within | A-Cov | D-Cov | Spanning | Within |
| NZ_CP007659.1 | NC_014925.1 | 36953 | 37046 | 319.63 | 906200 | 906301 | 89.36 | 6 | 17 | 18 | 100 | 100 | 100 | 100 |
| NZ_CP007659.1 | NC_014925.1 | 36953 | 37133 | 263.84 | 906200 | 906387 | 137.22 | 13 | 20 | 20 | 100 | 100 | 100 | 100 |
| NZ_CP007659.1 | NC_014925.1 | 36953 | 37152 | 254.4 | 906200 | 906409 | 135.31 | 9 | 19 | 20 | 100 | 100 | 100 | 100 |
| NZ_CP007659.1 | NC_014925.1 | 36999 | 37133 | 225.07 | 906256 | 906387 | 169.08 | 11 | 18 | 20 | 100 | 100 | 100 | 100 |
| NZ_CP007659.1 | NC_014925.1 | 36999 | 37152 | 217.61 | 906256 | 906409 | 161.88 | 7 | 18 | 20 | 100 | 100 | 100 | 100 |
| NZ_CP007659.1 | NC_014925.1 | 37045 | 37152 | 197.65 | 906300 | 906409 | 178.28 | 5 | 12 | 19 | 100 | 100 | 100 | 100 |
| NC_017763.1 | NC_002951.2 | 1554089 | 1562718 | 54.38 | 358442 | 369368 | 2.46 | 16 | 3 | 25 | 99 | 98 | 100 | 95 |
| NC_017763.1 | NC_002951.2 | 1554762 | 1562718 | 55.43 | 358442 | 369170 | 2.46 | 29 | 3 | 25 | 99 | 99 | 100 | 97 |
| NC_017763.1 | NC_014925.1 | 36952 | 37045 | 299.67 | 906200 | 906301 | 76.57 | 7 | 15 | 19 | 100 | 100 | 100 | 100 |
| NC_017763.1 | NC_014925.1 | 36952 | 37132 | 228.17 | 906200 | 906387 | 109.47 | 21 | 17 | 21 | 100 | 100 | 100 | 100 |
| NC_017763.1 | NC_014925.1 | 36952 | 37151 | 217.36 | 906200 | 906409 | 105.63 | 15 | 16 | 21 | 100 | 100 | 100 | 100 |
| NC_017763.1 | NC_014925.1 | 36998 | 37132 | 183.81 | 906256 | 906387 | 134.34 | 11 | 16 | 21 | 99 | 100 | 100 | 100 |
| NC_017763.1 | NC_014925.1 | 36998 | 37151 | 175.26 | 906256 | 906409 | 125.52 | 8 | 16 | 21 | 100 | 100 | 100 | 100 |
| NC_017763.1 | NC_014925.1 | 37044 | 37151 | 145.74 | 906300 | 906409 | 132.93 | 6 | 10 | 20 | 100 | 100 | 100 | 100 |
| NZ_CP007659.1 | NZ_CP011526.1 | 1568903 | 1575973 | 55.17 | 846399 | 854374 | 10.79 | 13 | 3 | 234 | 97 | 99 | 100 | 99 |
| NZ_CP007659.1 | NZ_CP011526.1 | 1568903 | 1579178 | 54.74 | 842251 | 854374 | 19.62 | 5 | 3 | 747 | 98 | 100 | 99 | 100 |
| NZ_CP007659.1 | NZ_CP011526.1 | 1568953 | 1575973 | 55.21 | 846399 | 854250 | 10.95 | 26 | 3 | 234 | 99 | 98 | 100 | 98 |
| NZ_CP007659.1 | NZ_CP011526.1 | 1568953 | 1579178 | 54.76 | 842251 | 854250 | 19.82 | 10 | 3 | 747 | 100 | 100 | 100 | 100 |
| NC_017763.1 | NZ_CP011526.1 | 1554717 | 1561787 | 55.17 | 846399 | 854374 | 10.79 | 13 | 3 | 234 | 99 | 100 | 100 | 100 |
| NC_017763.1 | NZ_CP011526.1 | 1554717 | 1564992 | 54.74 | 842251 | 854374 | 19.62 | 5 | 3 | 747 | 99 | 100 | 100 | 100 |
| NC_017763.1 | NZ_CP011526.1 | 1554767 | 1561787 | 55.21 | 846399 | 854250 | 10.95 | 26 | 3 | 234 | 99 | 100 | 100 | 100 |
| NC_017763.1 | NZ_CP011526.1 | 1554767 | 1564992 | 54.76 | 842251 | 854250 | 19.82 | 10 | 3 | 747 | 97 | 100 | 100 | 100 |
| NZ_CP007659.1 | NC_002951.2 | 1568275 | 1576904 | 54.38 | 358442 | 369368 | 2.46 | 16 | 3 | 25 | 99 | 98 | 100 | 96 |
| NZ_CP007659.1 | NC_002951.2 | 1568948 | 1576904 | 55.43 | 358442 | 369170 | 2.46 | 29 | 3 | 25 | 99 | 99 | 100 | 97 |

**Table S22:** Acceptor and donor candidates for ERR103395 run with yara, species filter and no samflag filter. Sampling sensitivity = 85. No taxon blacklist. No parent blacklist. No species blacklist. (-)0.000^*\**^represents absolute values 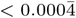.

| Type | Candidate |  | MicrobeGPS metrics |  |  | DaisyGPS metrics |  |
| --- | --- | --- | --- | --- | --- | --- | --- |
|  | Name | Accession.Version | Number Reads | Validity | Heterogeneity | Property | Property Score |
| Acceptor | Staphylococcus aureus subsp. aureus ECT-R 2 | NC_017343.1 | 120322 | 0.591 | 0.070 | 0.521 | 0.013 |
| Acceptor | Staphylococcus aureus subsp. aureus N315 | NC_002745.2 | 121110 | 0.576 | 0.069 | 0.507 | 0.013 |
| Donor | Enterococcus faecium Aus0004 | NC_017022.1 | 471 | 0.001 | 0.974 | -0.973 | -0.000* |
| Donor | Staphylococcus epidermidis ATCC 12228 | NC_004461.1 | 391 | 0.001 | 0.971 | -0.97 | -0.000* |
| Donor | Staphylococcus pseudintermedius HKU10-03 | NC_014925.1 | 470 | 0.001 | 0.806 | -0.805 | -0.000* |
| Donor | Staphylococcus lugdunensis HKU09-01 | NC_013893.1 | 59 | 0.003 | 0.765 | -0.762 | -0.000* |
| Donor | Staphylococcus warneri SG1 | NC_020164.1 | 294 | 0.011 | 0.693 | -0.683 | -0.000* |
| Donor | Staphylococcus haemolyticus JCSC1435 | NC_007168.1 | 362 | 0.002 | 0.556 | -0.554 | -0.000* |
| Acceptor-like Donor | Staphylococcus aureus subsp. aureus | NZ_CP009554.1 | 14824 | 0.093 | 0.091 | 0.002 | 0.000* |

**Table S23:** Results for ERR103395 run with yara, gustaf, species filter and no samflag filter. Sampling sensitivity = 90. Split read threshold = 3. No taxon blacklist. No parent blacklist. No species blacklist.

| Organism |  | Acceptor |  |  | Donor |  |  | Read Evidence |  |  | Evidence Filter |  |  |  |
| --- | --- | --- | --- | --- | --- | --- | --- | --- | --- | --- | --- | --- | --- | --- |
| Acceptor | Donor | Start | End | Coverage | Start | End | Coverage | Split | Spanning | Within | A-Cov | D-Cov | Spanning | Within |
| NC_002745.2 | NC_013893.1 | 2060607 | 2069048 | 16.44 | 2073055 | 2083555 | 11.08 | 28 | 8 | 339 | 1 | 100 | 100 | 100 |
| NC_002745.2 | NC_013893.1 | 2060607 | 2069067 | 16.47 | 2073055 | 2083576 | 11.07 | 18 | 6 | 339 | 2 | 100 | 99 | 100 |
| NC_002745.2 | NC_013893.1 | 2060762 | 2069048 | 16.74 | 2073192 | 2083555 | 11.11 | 7 | 8 | 339 | 1 | 100 | 100 | 100 |
| NC_002745.2 | NZ_CP009554.1 | 1142176 | 1142913 | 0.26 | 685582 | 686374 | 22.86 | 8 | 11 | 52 | 3 | 100 | 100 | 100 |
| NC_002745.2 | NZ_CP009554.1 | 1142176 | 1142913 | 0.26 | 685582 | 717267 | 0.66 | 12 | 14 | 53 | 5 | 94 | 97 | 94 |
| NC_002745.2 | NZ_CP009554.1 | 1142912 | 1142913 | 1.0 | 685581 | 716475 | 0.66 | 11 | 12 | 53 | 4 | 93 | 98 | 94 |
| NC_002745.2 | NZ_CP009554.1 | 2056699 | 2058174 | 0.02 | 2150234 | 2162636 | 2.65 | 10 | 6 | 43 | 2 | 98 | 99 | 94 |
| NC_002745.2 | NZ_CP009554.1 | 2056699 | 2060475 | 5.31 | 2150276 | 2162636 | 2.66 | 13 | 6 | 43 | 1 | 100 | 99 | 99 |
| NC_002745.2 | NZ_CP009554.1 | 2056699 | 2069076 | 10.31 | 2158985 | 2162636 | 3.14 | 24 | 34 | 8 | 2 | 99 | 100 | 90 |
| NC_002745.2 | NZ_CP009554.1 | 2058173 | 2069076 | 11.7 | 2150233 | 2158985 | 2.44 | 51 | 5 | 34 | 0 | 97 | 100 | 97 |
| NC_002745.2 | NZ_CP009554.1 | 2058173 | 2069105 | 11.7 | 2150233 | 2159011 | 2.48 | 7 | 5 | 34 | 3 | 100 | 100 | 98 |
| NC_002745.2 | NZ_CP009554.1 | 2058173 | 2069324 | 11.5 | 2150233 | 2159202 | 3.5 | 27 | 26 | 43 | 0 | 100 | 100 | 99 |
| NC_002745.2 | NZ_CP009554.1 | 2058173 | 2069355 | 11.57 | 2150233 | 2159253 | 3.62 | 7 | 26 | 43 | 0 | 100 | 100 | 96 |
| NC_002745.2 | NZ_CP009554.1 | 2060474 | 2069076 | 12.5 | 2150275 | 2158985 | 2.45 | 52 | 5 | 34 | 0 | 99 | 100 | 99 |
| NC_002745.2 | NZ_CP009554.1 | 2060474 | 2069105 | 12.5 | 2150275 | 2159011 | 2.49 | 8 | 5 | 34 | 2 | 100 | 100 | 98 |
| NC_002745.2 | NZ_CP009554.1 | 2060474 | 2069324 | 12.23 | 2150275 | 2159202 | 3.51 | 28 | 26 | 43 | 5 | 98 | 100 | 97 |
| NC_002745.2 | NZ_CP009554.1 | 2060474 | 2069355 | 12.31 | 2150275 | 2159253 | 3.63 | 8 | 26 | 43 | 1 | 100 | 100 | 99 |
| NC_002745.2 | NZ_CP009554.1 | 2060607 | 2065052 | 19.41 | 364874 | 369569 | 10.13 | 14 | 5 | 133 | 2 | 100 | 100 | 100 |
| NC_002745.2 | NZ_CP009554.1 | 2060607 | 2068738 | 12.04 | 361186 | 369569 | 7.19 | 18 | 1 | 154 | 0 | 100 | 100 | 100 |
| NC_002745.2 | NZ_CP009554.1 | 2065051 | 2068738 | 3.16 | 361186 | 364873 | 3.44 | 9 | 3 | 21 | 3 | 98 | 99 | 97 |
| NC_002745.2 | NZ_CP009554.1 | 2069075 | 2069324 | 2.76 | 2158984 | 2159202 | 45.94 | 89 | 32 | 8 | 4 | 100 | 100 | 100 |
| NC_002745.2 | NZ_CP009554.1 | 2069075 | 2069355 | 6.49 | 2158984 | 2159253 | 41.82 | 9 | 38 | 8 | 2 | 100 | 100 | 99 |
| NC_002745.2 | NZ_CP009554.1 | 2069104 | 2069324 | 1.65 | 2159010 | 2159202 | 50.14 | 12 | 32 | 8 | 3 | 100 | 100 | 99 |
| NC_017343.1 | NZ_CP009554.1 | 1100697 | 1101434 | 0.42 | 685582 | 686374 | 22.86 | 8 | 11 | 52 | 1 | 100 | 100 | 100 |
| NC_017343.1 | NZ_CP009554.1 | 1100697 | 1101434 | 0.42 | 685582 | 717267 | 0.66 | 15 | 14 | 53 | 1 | 99 | 99 | 98 |
| NC_017343.1 | NZ_CP009554.1 | 1101433 | 1101434 | 1.0 | 685581 | 716475 | 0.66 | 11 | 12 | 53 | 0 | 98 | 96 | 96 |
| NC_002745.2 | NC_004461.1 | 61651 | 61779 | 4.23 | 37793 | 55322 | 7.62 | 30 | 73 | 392 | 4 | 100 | 99 | 100 |
| NC_002745.2 | NC_004461.1 | 61651 | 61799 | 3.8 | 37814 | 55322 | 7.62 | 14 | 73 | 392 | 4 | 100 | 100 | 100 |
| NC_002745.2 | NC_004461.1 | 61651 | 61851 | 2.83 | 37866 | 55322 | 7.61 | 18 | 73 | 392 | 2 | 100 | 99 | 100 |
| NC_002745.2 | NC_004461.1 | 61755 | 61779 | 3.04 | 37793 | 55383 | 7.59 | 12 | 73 | 392 | 2 | 100 | 100 | 100 |
| NC_002745.2 | NC_004461.1 | 61755 | 61799 | 2.14 | 37814 | 55383 | 7.59 | 8 | 73 | 392 | 5 | 100 | 99 | 100 |
| NC_002745.2 | NC_004461.1 | 61755 | 61851 | 1.01 | 37866 | 55383 | 7.58 | 9 | 73 | 392 | 3 | 100 | 100 | 100 |
| NC_002745.2 | NC_004461.1 | 61778 | 62058 | 2.57 | 37792 | 57274 | 6.86 | 7 | 73 | 392 | 1 | 100 | 100 | 100 |
| NC_002745.2 | NC_004461.1 | 61778 | 62354 | 7.14 | 37792 | 57575 | 6.75 | 7 | 68 | 392 | 2 | 100 | 100 | 100 |
| NC_002745.2 | NC_004461.1 | 61778 | 62414 | 7.02 | 37792 | 57608 | 6.74 | 8 | 64 | 392 | 1 | 100 | 99 | 100 |
| NC_002745.2 | NC_004461.1 | 61798 | 62058 | 2.68 | 37813 | 57274 | 6.85 | 3 | 73 | 392 | 5 | 100 | 100 | 100 |
| NC_002745.2 | NC_004461.1 | 61798 | 62354 | 7.35 | 37813 | 57575 | 6.75 | 3 | 68 | 392 | 1 | 100 | 99 | 100 |
| NC_002745.2 | NC_004461.1 | 61798 | 62414 | 7.21 | 37813 | 57608 | 6.74 | 4 | 64 | 392 | 1 | 100 | 100 | 100 |
| NC_002745.2 | NC_004461.1 | 61850 | 62058 | 3.34 | 37865 | 57274 | 6.84 | 4 | 73 | 392 | 5 | 100 | 99 | 100 |
| NC_002745.2 | NC_004461.1 | 61850 | 62354 | 8.11 | 37865 | 57575 | 6.74 | 4 | 68 | 392 | 4 | 100 | 100 | 100 |
| NC_002745.2 | NC_004461.1 | 61850 | 62414 | 7.87 | 37865 | 57608 | 6.73 | 7 | 64 | 392 | 2 | 100 | 100 | 100 |

**Table S24:** Acceptor and donor candidates for ERR103396 run with yara, species filter and no samflag filter. Sampling sensitivity = 85. No taxon blacklist. No parent blacklist. No species blacklist. (-)0.000^*\**^represents absolute values 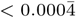.

| Candidate |  |  | MicrobeGPS metrics |  |  | DaisyGPS metrics |  |
| --- | --- | --- | --- | --- | --- | --- | --- |
| Type | Name | Accession.Version | Number Reads | Validity | Heterogeneity | Property | Property Score |
| Acceptor | Staphylococcus aureus subsp. aureus HO 5096 0412 | NC_017763.1 | 222016 | 0.817 | 0.042 | 0.775 | 0.043 |
| Acceptor | Staphylococcus aureus subsp. aureus | NZ_CP007659.1 | 223952 | 0.815 | 0.049 | 0.767 | 0.043 |
| Donor | Staphylococcus pseudintermedius ED99 | NC_017568.1 | 536 | 0.002 | 0.708 | -0.707 | -0.000* |
| Donor | Staphylococcus warneri SG1 | NC_020164.1 | 267 | 0.003 | 0.696 | -0.693 | -0.000* |
| Donor | Staphylococcus epidermidis RP62A | NC_002976.3 | 1067 | 0.003 | 0.582 | -0.579 | -0.000* |
| Donor | Staphylococcus haemolyticus JCSC1435 | NC_007168.1 | 370 | 0.003 | 0.492 | -0.489 | -0.000* |
| Donor | Staphylococcus aureus subsp. aureus COL | NC_002951.2 | 21752 | 0.098 | 0.156 | -0.058 | -0.000* |
| Acceptor-like Donor | Staphylococcus aureus subsp. aureus | NZ_CP012012.1 | 21332 | 0.097 | 0.094 | 0.003 | 0.000* |

**Table S25:** Results for ERR103396 run with yara, gustaf, species filter and no samflag filter. Sampling sensitivity = 90. Split read threshold = 3. No taxon blacklist. No parent blacklist. No species blacklist.

| Organism |  | Acceptor |  |  | Donor |  |  | Read Evidence |  |  | Evidence Filter |  |  |  |
| --- | --- | --- | --- | --- | --- | --- | --- | --- | --- | --- | --- | --- | --- | --- |
| Acceptor | Donor | Start | End | Coverage | Start | End | Coverage | Split | Spanning | Within | A-Cov | D-Cov | Spanning | Within |
| NC_017763.1 | NZ_CP012012.1 | 98589 | 98635 | 95.67 | 125862 | 126004 | 35.02 | 3 | 20 | 5 | 100 | 100 | 100 | 100 |
| NC_017763.1 | NC_017568.1 | 409730 | 409775 | 16.98 | 2481624 | 2485653 | 3.95 | 14 | 5 | 35 | 1 | 100 | 100 | 100 |

**Table S26:** Acceptor and donor candidates for ERR103397 run with yara, species filter and no samflag filter. Sampling sensitivity = 85. No taxon blacklist. No parent blacklist. No species blacklist. (-)0.000^*\**^represents absolute values 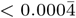.

| Candidate |  |  | MicrobeGPS metrics |  |  | DaisyGPS metrics |  |
| --- | --- | --- | --- | --- | --- | --- | --- |
| Type | Name | Accession.Version | Number Reads | Validity | Heterogeneity | Property | Property Score |
| Acceptor | Staphylococcus aureus subsp. aureus MSSA476 | NC_002953.3 | 84971 | 0.634 | 0.094 | 0.540 | 0.017 |
| Acceptor | Staphylococcus aureus subsp. aureus MW2 | NC_003923.1 | 83556 | 0.621 | 0.089 | 0.531 | 0.017 |
| Donor | Staphylococcus pseudintermedius HKU10-03 | NC_014925.1 | 3645 | 0.002 | 0.744 | -0.742 | -0.001 |
| Donor | Staphylococcus warneri SG1 | NC_020164.1 | 168 | 0.003 | 0.69 | -0.697 | -0.000* |
| Donor | Staphylococcus haemolyticus JCSC1435 | NC_007168.1 | 2650 | 0.004 | 0.604 | -0.600 | -0.001 |
| Donor | Staphylococcus epidermidis RP62A | NC_002976.3 | 1082 | 0.002 | 0.583 | -0.581 | -0.000* |
| Donor | Staphylococcus lugdunensis HKU09-01 | NC_013893.1 | 3709 | 0.004 | 0.356 | -0.352 | -0.001 |
| Donor | Staphylococcus aureus subsp. aureus | NZ_CP009554.1 | 19819 | 0.092 | 0.314 | -0.222 | -0.002 |
| Acceptor-like Donor | Staphylococcus aureus subsp. aureus | NZ_CP009361.1 | 9253 | 0.097 | 0.092 | 0.005 | 0.000* |

**Table S27:** Results for ERR103397 run with yara, gustaf, species filter and no samflag filter. Sampling sensitivity = 90. Split read threshold = 3. No taxon blacklist. No parent blacklist. No species blacklist.

| Organism |  | Acceptor |  |  | Donor |  |  | Read Evidence |  |  | Evidence Filter |  |  |  |
| --- | --- | --- | --- | --- | --- | --- | --- | --- | --- | --- | --- | --- | --- | --- |
| Acceptor | Donor | Start | End | Coverage | Start | End | Coverage | Split | Spanning | Within | A-Cov | D-Cov | Spanning | Within |
| NC_003923.1 | NC_007168.1 | 44986 | 45306 | 40.24 | 66689 | 67028 | 5.47 | 6 | 1 | 2 | 98 | 100 | 100 | 100 |
| NC_003923.1 | NC_002976.3 | 44776 | 44988 | 12.98 | 2520640 | 2520803 | 10.01 | 4 | 4 | 1 | 6 | 100 | 100 | 100 |
| NC_003923.1 | NC_002976.3 | 44987 | 45380 | 36.37 | 2520639 | 2561294 | 0.64 | 14 | 58 | 46 | 96 | 95 | 98 | 95 |
| NC_003923.1 | NC_002976.3 | 44987 | 45606 | 31.5 | 2520639 | 2561094 | 0.63 | 4 | 58 | 44 | 95 | 96 | 100 | 97 |
| NC_003923.1 | NC_002976.3 | 45026 | 45380 | 37.73 | 2561294 | 2561636 | 10.28 | 14 | 3 | 3 | 100 | 99 | 100 | 99 |
| NC_003923.1 | NC_002976.3 | 45026 | 45606 | 32.0 | 2561094 | 2561636 | 7.1 | 4 | 3 | 4 | 92 | 98 | 100 | 98 |
| NC_003923.1 | NC_002976.3 | 45026 | 45870 | 27.86 | 2560793 | 2561636 | 6.17 | 6 | 4 | 4 | 91 | 99 | 100 | 100 |
| NC_003923.1 | NC_002976.3 | 45070 | 45307 | 38.1 | 2561337 | 2561580 | 12.23 | 4 | 5 | 3 | 94 | 100 | 100 | 100 |
| NC_003923.1 | NC_002976.3 | 45070 | 45380 | 38.28 | 2561294 | 2561580 | 12.12 | 40 | 5 | 3 | 98 | 100 | 100 | 100 |
| NC_002953.3 | NC_007168.1 | 41508 | 57483 | 26.01 | 67036 | 120082 | 2.7 | 20 | 4 | 471 | 94 | 98 | 95 | 98 |

**Table S28:** Acceptor and donor candidates for ERR103398 run with yara, species filter and no samflag filter. Sampling sensitivity = 85. No taxon blacklist. No parent blacklist. No species blacklist. (-)0.000^*\**^represents absolute values 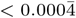.

| Candidate |  |  | MicrobeGPS metrics |  |  | DaisyGPS metrics |  |
| --- | --- | --- | --- | --- | --- | --- | --- |
| Type | Name | Accession.Version | Number Reads | Validity | Heterogeneity | Property | Property Score |
| Acceptor | Staphylococcus aureus subsp. aureus MSSA476 | NC_002953.3 | 192949 | 0.671 | 0.11 | 0.562 | 0.017 |
| Acceptor | Staphylococcus aureus subsp. aureus MW2 | NC_003923.1 | 189418 | 0.658 | 0.103 | 0.555 | 0.016 |
| Donor | Staphylococcus pseudintermedius HKU10-03 | NC_014925.1 | 16866 | 0.002 | 0.745 | -0.742 | -0.002 |
| Donor | Staphylococcus warneri SG1 | NC_020164.1 | 461 | 0.003 | 0.69 | -0.697 | -0.000* |
| Donor | Staphylococcus epidermidis PM221 | NZ_HG813242.1 | 4779 | 0.001 | 0.656 | -0.655 | -0.001 |
| Donor | Staphylococcus haemolyticus JCSC1435 | NC_007168.1 | 12023 | 0.004 | 0.636 | -0.632 | -0.001 |
| Donor | Staphylococcus lugdunensis HKU09-01 | NC_013893.1 | 16800 | 0.004 | 0.356 | -0.351 | -0.001 |
| Donor | Staphylococcus aureus subsp. aureus | NZ_CP009554.1 | 70966 | 0.095 | 0.398 | -0.304 | -0.003 |
| Acceptor-like Donor | Staphylococcus aureus CA-347 | NC_021554.1 | 18666 | 0.098 | 0.090 | 0.007 | 0.000* |

**Table S29:** Results for ERR103398 run with yara, gustaf, species filter and no samflag filter. Sampling sensitivity = 90. Split read threshold = 3. No taxon blacklist. No parent blacklist. No species blacklist.

| Organism |  | Acceptor |  |  | Donor |  |  | Read Evidence |  |  | Evidence Filter |  |  |  |
| --- | --- | --- | --- | --- | --- | --- | --- | --- | --- | --- | --- | --- | --- | --- |
| Acceptor | Donor | Start | End | Coverage | Start | End | Coverage | Split | Spanning | Within | A-Cov | D-Cov | Spanning | Within |
| NC_002953.3 | NC_007168.1 | 27843 | 41291 | 63.48 | 67243 | 97433 | 4.35 | 18 | 2 | 436 | 95 | 97 | 99 | 99 |
| NC_002953.3 | NC_007168.1 | 27843 | 41907 | 63.94 | 66699 | 97433 | 4.54 | 3 | 4 | 446 | 91 | 99 | 99 | 99 |
| NC_002953.3 | NC_007168.1 | 27843 | 57484 | 69.06 | 94688 | 97433 | 47.35 | 7 | 2 | 431 | 99 | 100 | 100 | 100 |
| NC_002953.3 | NC_007168.1 | 41290 | 41907 | 73.94 | 66699 | 67242 | 15.3 | 9 | 6 | 14 | 96 | 100 | 100 | 100 |
| NC_002953.3 | NC_007168.1 | 41290 | 57483 | 73.69 | 67242 | 120082 | 6.24 | 77 | 6 | 1105 | 99 | 94 | 99 | 97 |
| NC_002953.3 | NC_007168.1 | 41290 | 57484 | 73.69 | 30072 | 67242 | 0.28 | 25 | 7 | 16 | 100 | 93 | 100 | 97 |
| NC_002953.3 | NC_007168.1 | 41508 | 41605 | 95.51 | 66925 | 67036 | 6.53 | 17 | 6 | 1 | 97 | 100 | 100 | 100 |
| NC_002953.3 | NC_007168.1 | 41508 | 41907 | 90.78 | 66699 | 67036 | 7.76 | 5 | 1 | 2 | 98 | 100 | 100 | 100 |
| NC_002953.3 | NC_007168.1 | 41508 | 57483 | 74.11 | 67036 | 120082 | 6.32 | 39 | 10 | 1111 | 98 | 91 | 94 | 95 |
| NC_002953.3 | NC_007168.1 | 41508 | 57484 | 74.11 | 67036 | 94688 | 0.25 | 13 | 8 | 11 | 100 | 93 | 97 | 95 |
| NC_002953.3 | NC_007168.1 | 41604 | 41907 | 89.3 | 66699 | 66924 | 8.38 | 9 | 2 | 1 | 97 | 99 | 100 | 99 |
| NC_002953.3 | NC_007168.1 | 41604 | 57483 | 73.98 | 66924 | 120082 | 6.32 | 77 | 15 | 1112 | 100 | 92 | 99 | 94 |
| NC_002953.3 | NC_007168.1 | 41906 | 57483 | 73.68 | 66698 | 120082 | 6.33 | 21 | 15 | 1115 | 100 | 91 | 100 | 94 |
| NC_002953.3 | NC_007168.1 | 41906 | 57484 | 73.68 | 66698 | 94688 | 0.34 | 8 | 13 | 15 | 100 | 92 | 100 | 96 |
| NC_002953.3 | NZ_HG813242.1 | 34149 | 41829 | 68.51 | 35795 | 85587 | 4.98 | 78 | 19 | 287 | 99 | 99 | 97 | 99 |
| NC_002953.3 | NZ_HG813242.1 | 34149 | 41907 | 68.57 | 35721 | 85587 | 4.97 | 50 | 19 | 287 | 95 | 97 | 98 | 97 |
| NC_002953.3 | NZ_HG813242.1 | 34149 | 57484 | 71.98 | 54292 | 85587 | 6.1 | 34 | 3 | 280 | 97 | 97 | 97 | 97 |
| NC_002953.3 | NZ_HG813242.1 | 34149 | 57484 | 71.98 | 57395 | 85587 | 5.91 | 42 | 2 | 207 | 100 | 97 | 100 | 97 |
| NC_002953.3 | NZ_HG813242.1 | 34180 | 41829 | 68.6 | 35795 | 85647 | 4.97 | 240 | 19 | 287 | 93 | 96 | 97 | 96 |
| NC_002953.3 | NZ_HG813242.1 | 34180 | 41907 | 68.66 | 35721 | 85647 | 4.96 | 142 | 19 | 287 | 97 | 95 | 96 | 95 |
| NC_002953.3 | NZ_HG813242.1 | 34180 | 57484 | 72.02 | 54292 | 85647 | 6.09 | 86 | 3 | 280 | 100 | 96 | 100 | 96 |
| NC_002953.3 | NZ_HG813242.1 | 34180 | 57484 | 72.02 | 57395 | 85647 | 5.9 | 114 | 2 | 207 | 100 | 97 | 100 | 97 |
| NC_002953.3 | NZ_HG813242.1 | 41828 | 56986 | 72.4 | 35794 | 85689 | 4.98 | 81 | 19 | 287 | 99 | 94 | 97 | 94 |
| NC_002953.3 | NZ_HG813242.1 | 41828 | 57484 | 73.68 | 35794 | 57395 | 3.75 | 43 | 2 | 70 | 99 | 94 | 100 | 93 |
| NC_002953.3 | NZ_HG813242.1 | 41828 | 57484 | 73.68 | 35794 | 62757 | 9.18 | 15 | 3 | 286 | 100 | 97 | 97 | 97 |
| NC_002953.3 | NZ_HG813242.1 | 41906 | 56986 | 72.39 | 35720 | 85689 | 4.97 | 143 | 19 | 287 | 100 | 93 | 98 | 93 |
| NC_002953.3 | NZ_HG813242.1 | 41906 | 57484 | 73.68 | 35720 | 57395 | 3.74 | 67 | 2 | 70 | 100 | 98 | 98 | 98 |
| NC_002953.3 | NZ_HG813242.1 | 41906 | 57484 | 73.68 | 35720 | 62757 | 9.16 | 11 | 3 | 286 | 100 | 99 | 98 | 99 |
| NC_002953.3 | NZ_HG813242.1 | 56985 | 57484 | 112.78 | 54292 | 85688 | 6.1 | 64 | 3 | 280 | 100 | 97 | 98 | 97 |
| NC_002953.3 | NZ_HG813242.1 | 56985 | 57484 | 112.78 | 57395 | 85688 | 5.91 | 80 | 2 | 207 | 100 | 98 | 99 | 98 |
| NC_003923.1 | NC_007168.1 | 44606 | 45306 | 69.87 | 66689 | 67411 | 10.45 | 29 | 2 | 16 | 95 | 100 | 100 | 100 |
| NC_003923.1 | NC_007168.1 | 45026 | 45143 | 97.32 | 66870 | 66987 | 2.7 | 9 | 4 | 2 | 100 | 99 | 100 | 99 |
| NC_003923.1 | NC_007168.1 | 45026 | 45306 | 93.51 | 66689 | 66987 | 17.21 | 25 | 6 | 13 | 98 | 100 | 100 | 100 |
| NC_003923.1 | NC_007168.1 | 45062 | 45306 | 93.32 | 66689 | 66930 | 20.85 | 19 | 3 | 11 | 99 | 99 | 100 | 100 |
| NC_003923.1 | NC_007168.1 | 45082 | 45306 | 92.39 | 66689 | 66929 | 20.93 | 10 | 4 | 11 | 98 | 100 | 100 | 100 |
| NC_003923.1 | NC_007168.1 | 45142 | 45306 | 90.77 | 66689 | 66869 | 26.71 | 10 | 5 | 11 | 97 | 99 | 100 | 99 |
| NC_002953.3 | NC_021554.1 | 41508 | 41593 | 91.24 | 60998 | 61108 | 10.47 | 19 | 2 | 5 | 96 | 99 | 100 | 99 |
| NC_002953.3 | NC_021554.1 | 41508 | 41884 | 81.12 | 60998 | 61399 | 20.48 | 7 | 1 | 11 | 95 | 99 | 100 | 99 |
| NC_002953.3 | NC_021554.1 | 41548 | 41884 | 80.93 | 61045 | 61399 | 21.66 | 4 | 2 | 9 | 96 | 100 | 100 | 100 |
| NC_002953.3 | NC_021554.1 | 41592 | 41884 | 78.23 | 61107 | 61399 | 24.23 | 22 | 3 | 8 | 92 | 99 | 100 | 99 |
| NC_003923.1 | NC_021554.1 | 44986 | 45384 | 76.26 | 61006 | 61391 | 27.57 | 55 | 2 | 20 | 90 | 100 | 100 | 100 |
| NC_003923.1 | NC_021554.1 | 45026 | 45306 | 80.71 | 61045 | 61347 | 29.43 | 7 | 3 | 18 | 95 | 100 | 100 | 100 |
| NC_003923.1 | NC_021554.1 | 45026 | 45384 | 76.52 | 61045 | 61391 | 29.23 | 113 | 3 | 18 | 94 | 100 | 100 | 100 |
| NC_003923.1 | NC_021554.1 | 45062 | 45306 | 78.63 | 61102 | 61347 | 35.75 | 3 | 1 | 14 | 92 | 100 | 100 | 99 |
| NC_003923.1 | NC_021554.1 | 45062 | 45384 | 74.48 | 61102 | 61391 | 34.55 | 109 | 1 | 14 | 92 | 100 | 100 | 100 |
| NC_003923.1 | NC_021554.1 | 45082 | 45384 | 72.6 | 61105 | 61391 | 34.9 | 55 | 1 | 14 | 91 | 100 | 100 | 100 |

**Table S30:**
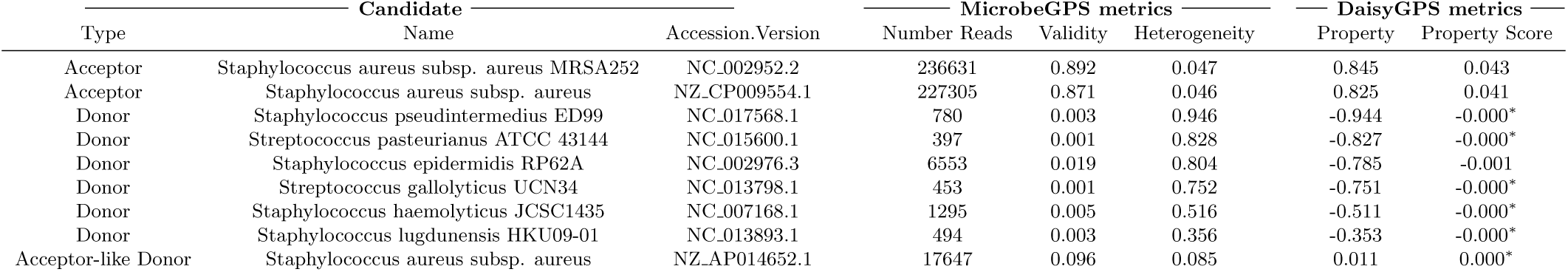
Acceptor and donor candidates for ERR159680 run with yara, species filter and no samflag filter. Sampling sensitivity = 85. No taxon blacklist. No parent blacklist. No species blacklist. (-)0.000^*\**^represents absolute values 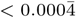.

| Type | Candidate |  | Accession.Version | MicrobeGPS metrics |  |  | DaisyGPS metrics |  |
| --- | --- | --- | --- | --- | --- | --- | --- | --- |
|  | Name |  |  | Number Reads | Validity | Heterogeneity | Property | Property Score |
| Acceptor | Staphylococcus aureus subsp. aureus MRSA252 |  | NC_002952.2 | 236631 | 0.892 | 0.047 | 0.845 | 0.043 |
| Acceptor | Staphylococcus aureus subsp. aureus |  | NZ_CP009554.1 | 227305 | 0.871 | 0.046 | 0.825 | 0.041 |
| Donor | Staphylococcus pseudintermedius ED99 |  | NC_017568.1 | 780 | 0.003 | 0.946 | -0.944 | -0.000* |
| Donor | Streptococcus pasteurianus ATCC 43144 |  | NC_015600.1 | 397 | 0.001 | 0.828 | -0.827 | -0.000* |
| Donor | Staphylococcus epidermidis RP62A |  | NC_002976.3 | 6553 | 0.019 | 0.804 | -0.785 | -0.001 |
| Donor | Streptococcus gallolyticus UCN34 |  | NC_013798.1 | 453 | 0.001 | 0.752 | -0.751 | -0.000* |
| Donor | Staphylococcus haemolyticus JCSC1435 |  | NC_007168.1 | 1295 | 0.005 | 0.516 | -0.511 | -0.000* |
| Donor | Staphylococcus lugdunensis HKU09-01 |  | NC_013893.1 | 494 | 0.003 | 0.356 | -0.353 | -0.000* |
| Acceptor-like Donor | Staphylococcus aureus subsp. aureus |  | NZ_AP014652.1 | 17647 | 0.096 | 0.085 | 0.011 | 0.000* |

**Table S31:** Results for ERR159680 run with yara, gustaf, species filter and no samflag filter. Sampling sensitivity = 90. Split read threshold = 3. No taxon blacklist. No parent blacklist. No species blacklist.

| Organism |  | Acceptor |  |  | Donor |  |  | Read Evidence |  |  | Evidence Filter |  |  |  |
| --- | --- | --- | --- | --- | --- | --- | --- | --- | --- | --- | --- | --- | --- | --- |
| Acceptor | Donor | Start | End | Coverage | Start | End | Coverage | Split | Spanning | Within | A-Cov | D-Cov | Spanning | Within |
| NZ_CP009554.1 | NC_002976.3 | 34120 | 34123 | 12.67 | 2536574 | 2584194 | 38.98 | 30 | 19 | 5752 | 6 | 100 | 98 | 100 |
| NZ_CP009554.1 | NC_002976.3 | 859613 | 866305 | 63.76 | 1398260 | 1404973 | 0.3 | 27 | 4 | 5 | 100 | 98 | 100 | 98 |
| NZ_CP009554.1 | NC_013893.1 | 2130925 | 2133716 | 7.9 | 2343346 | 2345047 | 5.05 | 17 | 1 | 10 | 3 | 100 | 100 | 100 |
| NZ_CP009554.1 | NC_013893.1 | 2131388 | 2133716 | 4.04 | 2343670 | 2345047 | 6.23 | 16 | 1 | 10 | 0 | 100 | 100 | 100 |
| NC_002952.2 | NC_002976.3 | 906791 | 906792 | 12.0 | 1398259 | 1404972 | 0.3 | 18 | 4 | 5 | 1 | 94 | 100 | 97 |
| NZ_CP009554.1 | NZ_AP014652.1 | 414814 | 417301 | 46.9 | 438237 | 438358 | 11.31 | 4 | 2 | 5 | 96 | 99 | 100 | 99 |
| NZ_CP009554.1 | NZ_AP014652.1 | 2110903 | 2123964 | 16.67 | 2007772 | 2020977 | 17.83 | 6 | 4 | 762 | 0 | 98 | 99 | 98 |
| NZ_CP009554.1 | NZ_AP014652.1 | 2110903 | 2131317 | 11.83 | 2007772 | 2029781 | 21.15 | 3 | 4 | 1493 | 0 | 100 | 97 | 100 |
| NZ_CP009554.1 | NZ_AP014652.1 | 2110903 | 2134197 | 10.62 | 2007772 | 2030266 | 20.77 | 25 | 3 | 1493 | 0 | 99 | 99 | 99 |
| NZ_CP009554.1 | NZ_AP014652.1 | 2123963 | 2131021 | 1.86 | 2020976 | 2029466 | 27.01 | 5 | 1 | 729 | 0 | 98 | 100 | 98 |
| NZ_CP009554.1 | NZ_AP014652.1 | 2123963 | 2131317 | 3.23 | 2020976 | 2029781 | 26.14 | 7 | 2 | 731 | 1 | 98 | 100 | 98 |
| NZ_CP009554.1 | NZ_AP014652.1 | 2123963 | 2134197 | 2.91 | 2020976 | 2030266 | 24.94 | 51 | 1 | 731 | 0 | 100 | 99 | 100 |
| NZ_CP009554.1 | NZ_AP014652.1 | 2125253 | 2131317 | 1.84 | 2022297 | 2029781 | 30.59 | 3 | 3 | 731 | 0 | 98 | 100 | 98 |
| NZ_CP009554.1 | NZ_AP014652.1 | 2125253 | 2134197 | 1.92 | 2022297 | 2030266 | 28.92 | 25 | 2 | 731 | 0 | 99 | 100 | 99 |
| NZ_CP009554.1 | NZ_AP014652.1 | 2131020 | 2134197 | 5.24 | 2029465 | 2030266 | 2.97 | 25 | 1 | 2 | 2 | 97 | 100 | 97 |
| NZ_CP009554.1 | NZ_AP014652.1 | 2131316 | 2134197 | 2.09 | 2029780 | 2030266 | 3.19 | 49 | 6 | 2 | 1 | 97 | 100 | 96 |
| NC_002952.2 | NC_013893.1 | 413772 | 417366 | 53.37 | 2079996 | 2083590 | 0.78 | 5 | 4 | 2 | 100 | 99 | 100 | 100 |

**Table S32:** Acceptor and donor candidates for ERR103400 run with yara, species filter and no samflag filter. Sampling sensitivity = 85. No taxon blacklist. No parent blacklist. No species blacklist. (-)0.000^*\**^represents absolute values 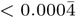.

| Type | Candidate |  | Accession.Version | MicrobeGPS metrics |  |  | DaisyGPS metrics |  |
| --- | --- | --- | --- | --- | --- | --- | --- | --- |
|  | Name |  |  | Number Reads | Validity | Heterogeneity | Property | Property Score |
| Acceptor | Staphylococcus aureus subsp. aureus HO 5096 0412 |  | NC_017763.1 | 484936 | 0.835 | 0.037 | 0.798 | 0.041 |
| Acceptor | Staphylococcus aureus subsp. aureus |  | NZ_CP007659.1 | 489699 | 0.832 | 0.048 | 0.784 | 0.041 |
| Donor | Staphylococcus haemolyticus JCSC1435 |  | NC_007168.1 | 3222 | 0.006 | 0.799 | -0.792 | -0.000* |
| Donor | Staphylococcus pseudintermedius ED99 |  | NC_017568.1 | 1398 | 0.002 | 0.701 | -0.699 | -0.000* |
| Donor | Staphylococcus warneri SG1 |  | NC_020164.1 | 583 | 0.003 | 0.695 | -0.692 | -0.000* |
| Donor | Staphylococcus epidermidis ATCC 12228 |  | NC_004461.1 | 3245 | 0.005 | 0.483 | -0.479 | -0.000* |
| Donor | Staphylococcus lugdunensis HKU09-01 |  | NC_013893.1 | 69 | 0.005 | 0.342 | -0.337 | -0.000* |
| Donor | Staphylococcus aureus subsp. aureus |  | NZ_CP009554.1 | 132861 | 0.21 | 0.254 | -0.044 | -0.001 |
| Acceptor-like Donor | Staphylococcus aureus subsp. aureus T0131 |  | NC_017347.1 | 50347 | 0.104 | 0.103 | 0.001 | 0.000* |

**Table S33:**
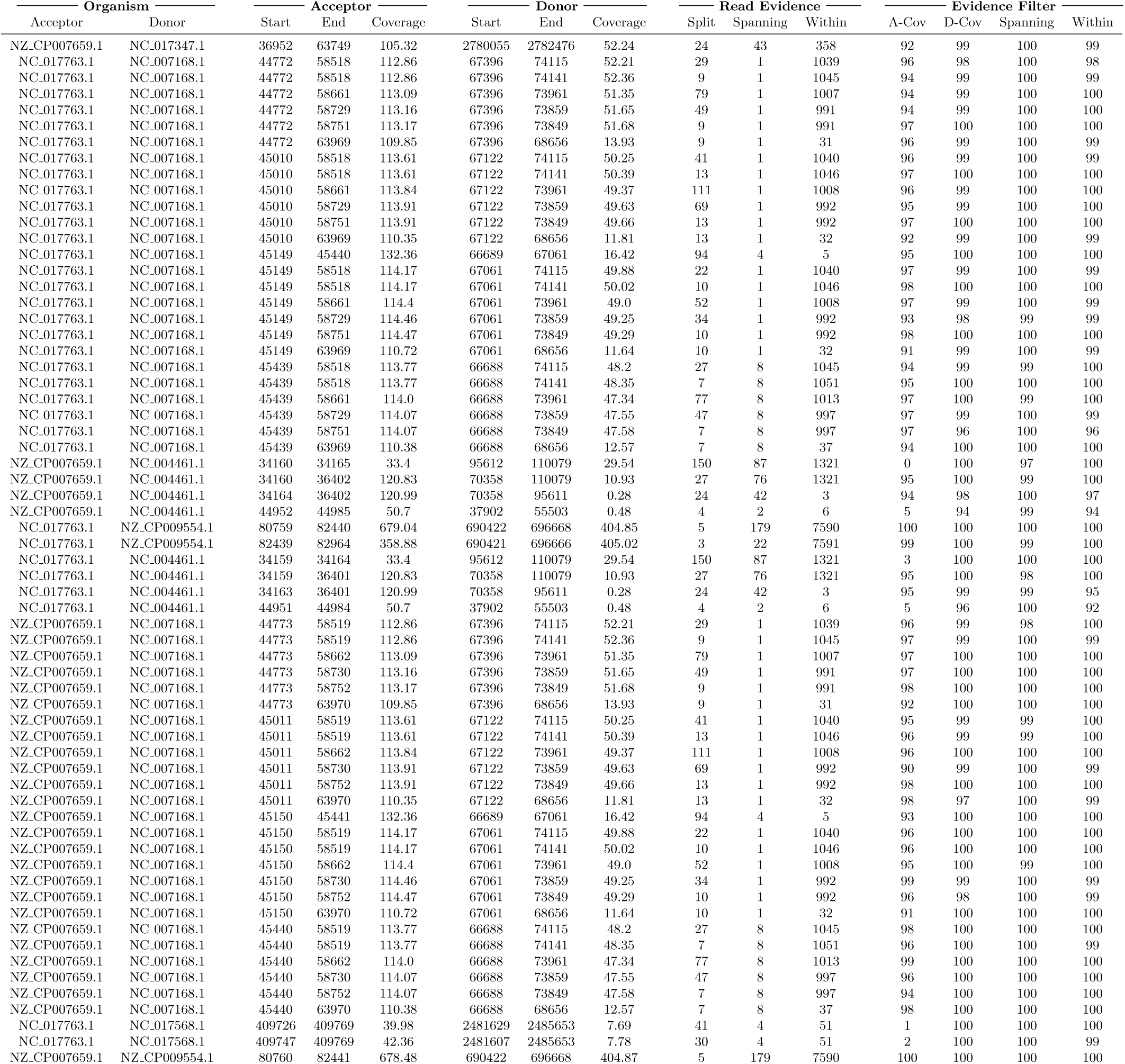
Results for ERR103400 run with yara, gustaf, species filter and no samflag filter. Sampling sensitivity = 90. Split read threshold = 3. No taxon blacklist. No parent blacklist. No species blacklist.

**Table S34:**
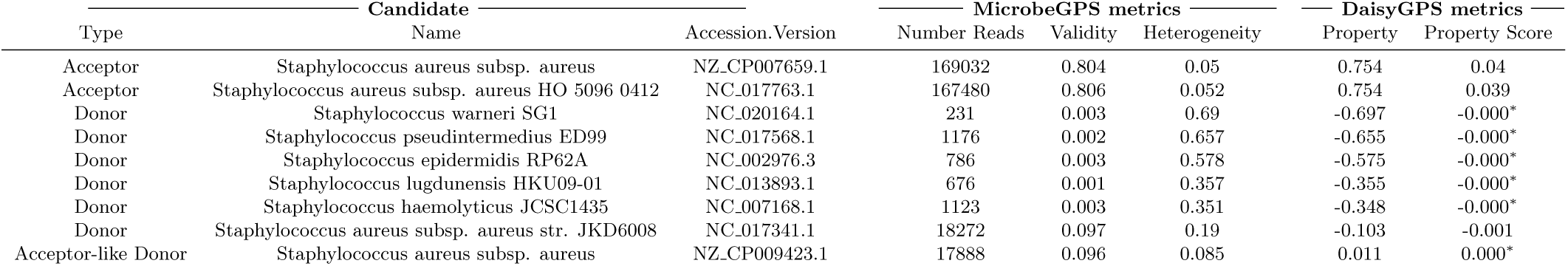
Acceptor and donor candidates for ERR103402 run with yara, species filter and no samflag filter. Sampling sensitivity = 85. No taxon blacklist. No parent blacklist. No species blacklist. (-)0.000^*\**^represents absolute values 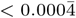.

**Table S35:** Results for ERR103402 run with yara, gustaf, species filter and no samflag filter. Sampling sensitivity = 90. Split read threshold = 3. No taxon blacklist. No parent blacklist. No species blacklist.

| Organism |  | Acceptor |  |  | Donor |  |  | Read Evidence |  |  | Evidence Filter |  |  |  |
| --- | --- | --- | --- | --- | --- | --- | --- | --- | --- | --- | --- | --- | --- | --- |
| Acceptor | Donor | Start | End | Coverage | Start | End | Coverage | Split | Spanning | Within | A-Cov | D-Cov | Spanning | Within |
| NZ_CP007659.1 | NC_020164.1 | 2038921 | 2038922 | 83.0 | 121511 | 123832 | 2.22 | 48 | 24 | 12 | 100 | 100 | 100 | 100 |
| NC_017763.1 | NC_020164.1 | 2024903 | 2024904 | 83.0 | 121511 | 123832 | 2.22 | 52 | 24 | 12 | 99 | 100 | 99 | 100 |
| NZ_CP007659.1 | NC_017341.1 | 2036785 | 2038062 | 62.16 | 2760130 | 2761402 | 144.91 | 6 | 48 | 540 | 99 | 100 | 100 | 100 |
| NZ_CP007659.1 | NC_013893.1 | 2036709 | 2038062 | 66.05 | 1722127 | 1723472 | 67.35 | 10 | 10 | 2 | 100 | 100 | 100 | 100 |
| NZ_CP007659.1 | NC_013893.1 | 2036785 | 2038063 | 62.02 | 949399 | 950670 | 74.87 | 9 | 1 | 3 | 99 | 100 | 100 | 100 |
| NZ_CP007659.1 | NC_013893.1 | 2036785 | 2038062 | 62.03 | 1722127 | 1723395 | 70.11 | 18 | 51 | 2 | 100 | 100 | 100 | 100 |
| NC_017763.1 | NC_013893.1 | 2022691 | 2024044 | 66.05 | 1722127 | 1723472 | 67.46 | 10 | 10 | 2 | 100 | 100 | 100 | 100 |
| NC_017763.1 | NC_013893.1 | 2022767 | 2024045 | 62.02 | 949399 | 950670 | 74.75 | 9 | 1 | 3 | 99 | 100 | 100 | 100 |
| NC_017763.1 | NC_013893.1 | 2022767 | 2024044 | 62.03 | 1722127 | 1723395 | 70.23 | 18 | 51 | 2 | 100 | 100 | 100 | 100 |
| NC_017763.1 | NC_017341.1 | 2022767 | 2024044 | 62.16 | 2760130 | 2761402 | 144.91 | 6 | 48 | 540 | 99 | 100 | 100 | 100 |
| NC_017763.1 | NC_007168.1 | 2022767 | 2024044 | 62.16 | 1828214 | 1829486 | 144.91 | 10 | 48 | 540 | 99 | 100 | 100 | 100 |
| NZ_CP007659.1 | NC_007168.1 | 2036785 | 2038062 | 62.16 | 1828214 | 1829486 | 144.91 | 10 | 48 | 540 | 100 | 100 | 100 | 100 |

